# HpBoRB, a helminth-derived CCP domain protein which binds RELMβ

**DOI:** 10.1101/2025.05.27.656348

**Authors:** Vivien Shek, Abhishek Jamwal, Danielle J Smyth, Tania Frangova, Alice R Savage, Sarah Kelly, Gavin J Wright, Rachel Toth, Rick M Maizels, Matthew K Higgins, Alasdair C Ivens, Hermelijn H Smits, Henry J McSorley

**Affiliations:** Division of Cell Signalling and Immunology, School of Life Sciences, University of Dundee, Dundee, United Kingdom; Department of Biochemistry, University of Oxford, Oxford, United Kingdom; Department of Biology, Hull York Medical School, York Biomedical Research Institute, University of York, York, United Kingdom; MRC Protein Phosphorylation and Ubiquitylation Unit, School of Life Sciences, University of Dundee, Dundee DD1 5EH, UK; Centre for Parasitology, School of Infection and Immunity, University of Glasgow, University, Glasgow, G12 8TA, UK; Institute of Immunology and Infection Research, School of Biological Sciences, University of Edinburgh, Edinburgh EH9 3JT, United Kingdom; Department of Immunology, Leiden University Medical Center, Leiden, Netherlands

**Keywords:** *Heligmosomoides polygyrus bakeri*, Complement Control Protein, immunomodulation, AVEXIS, protein-protein interaction, RELMβ

## Abstract

Helminth infections persist by influencing host immunity through the release of immunomodulatory proteins which prevent immune ejection. The intestinal nematode *Heligmosomoides polygyrus bakeri* (Hpb) secretes multiple families of immunomodulatory proteins, many of which are composed of consecutive Complement Control Protein (CCP) domains. We hypothesized that further CCP domain proteins are secreted by the parasite to interact with the host. We identified an unusually large number of CCP domain-containing proteins in the genome of Hpb, and cloned a range of these for screening in an Avidity-based Extracellular Interaction Screening (AVEXIS) assay, focussing on interactions with host immune proteins. This screen confirmed the binding of known immunomodulators (HpBARI, TGM1) for their targets (ST2, TGFBR2) and identified a new interaction between a 2 CCP domain Hpb protein and mouse resistin-like molecule beta (RELMβ), a host protein demonstrated to have anti-helminth properties. This protein was named Binder of RELMβ (HpBoRB). This interaction was confirmed in ELISA, competition assays, size exclusion chromatography and surface plasmon resonance experiments, identifying a subnanomolar affinity interaction between HpBoRB and RELMβ. These data may indicate that Hpb interferes with the potent anti-helminth host protein RELMβ and adds to our knowledge of the host-parasite interactions mediated by Hpb secreted proteins.

## 1. Introduction

Parasitic helminths are in an evolutionary arms race with their hosts. The host immune system has developed pathways for parasite ejection, while the parasite has co-evolved with its host to produce sophisticated modulators of the anti-parasite response (Fumagalli et al., 2011; Maizels et al., 2018; Ryan et al., 2022; Colomb and McSorley, 2025). This interaction allows many parasites to form chronic infections in their hosts, inhibiting type 2 immune responses against them. As the immune system evolved in the context of constant parasite infection and consequent immunosuppression, the immune system is prone to hyperactive type 2 immune responses in the absence of regular parasite infections, resulting in an epidemic of allergic disease in the developed world (Lambrecht and Hammad, 2017).

In recent years, some of the molecular basis of this interaction has become clear, as multiple parasite-derived immunomodulatory proteins have been identified which modulate the host immune response to the parasite’s advantage (Everts et al., 2012; Navarro et al., 2016; Johnston et al., 2017; Osbourn et al., 2017; de Los Reyes Jimenez et al., 2020; Vacca et al., 2020). These proteins are of interest due to their potential as treatments for allergic diseases such as asthma, and as vaccine candidates to prevent parasitic infections (Nisbet et al., 2013; Bancroft et al., 2019; Berkachy et al., 2021; Smyth et al., 2025).

One of the parasitic nematodes that has received the most interest as a modulator of host immunity is *Heligmosomoides polygyrus bakeri* (Hpb). The secretions of Hpb contain multiple families of immunoregulatory proteins including the HpARIs, which bind to IL-33 (Osbourn et al., 2017; Colomb et al., 2024a; Colomb et al., 2024b; Jamwal et al., 2024), the HpBARIs, which bind to the IL-33 receptor ST2 (Vacca et al., 2020), and the TGMs, which bind to the TGFβ receptor and cell-specific coreceptors (Johnston et al., 2017; Smyth et al., 2018; White et al., 2021; van Dinther et al., 2023; Maizels et al., 2025; Singh et al., 2025; White et al., 2025). These 3 families of proteins each contain 3-10 family members (3 HpARIs, 3 HpBARIs and 10 TGMs), and are structurally related to each other: each contain an N-terminal signal peptide followed by a string of consecutive atypical and nonidentical complement control protein (CCP) domains. CCP domains are present in many host and parasite proteins and are characterized by conserved sequence and structural elements, including 4 conserved cysteines per CCP domain. These conserved cysteines form disulphide bonds to stabilise the globular structure of the domain, consisting of 3 to 4 β-sheets (Jamwal et al., 2024).

We hypothesized that further host-parasite interactions could be mediated by Hpb CCP domain-containing proteins, therefore we assembled a complete list of proteins containing these domains in Hpb and other parasitic and free-living helminths, and confirmed that Hpb has dramatically expanded a family of CCP domain-containing proteins. We expressed a range of these Hpb CCP domain proteins and used an avidity-based extracellular interaction screening (AVEXIS) assay to identify Binder of RELMβ (HpBoRB), a secreted protein consisting of 2 CCP domains, which binds to host RELMβ. RELMβ is an effector molecule involved in the ejection of helminth parasites, and mice deficient in RELMβ are more susceptible to several intestinal nematodes, including Hpb (Herbert et al., 2009). Therefore, HpBoRB may represent part of the immunomodulatory armoury of Hpb.

## 2. Material and methods

### 2.1. Helminth genome analysis for secretory CCP domain superfamily proteins

All helminth genomic data were retrieved from WormBase ParaSite (Howe et al., 2017). The data mining tool BioMart was used to identify protein-coding genes containing CCP domain superfamily proteins using the Interpro domain IPR035976 (Blum et al., 2025) and signal peptides were identified by SignalP-5.0 (Almagro Armenteros et al., 2019).

### 2.2. H. polygyrus bakeri transcriptome

Adult *H. polygyrus bakeri* transcriptome data (produced in-house) was also used for analysis. In brief, purified adult worm mRNA was reverse transcribed to approximately 460,000 cDNA sequences. Sequencing was performed on a Roche 454 instrument resulting in reads of ∼200 nucleotides as described in (Hewitson et al., 2011).

### 2.3. *H. polygyrus bakeri* and murine immune recombinant protein expression

The signal peptides of selected protein candidates were identified using predictions from SignalP-5.0 (Almagro Armenteros et al., 2019) and excluded from subsequent gene synthesis. All AVEXIS bait and prey constructs were cloned from Hpb or mouse cDNA libraries where possible, or gene synthesised (GeneArt) if unavailable. Corresponding codon-optimised cDNA for human cell expression were synthesised with unique 5’*NotI* and 3’*AscI* restriction sites and subcloned into an expression plasmid. The plasmids used in this project are: AVEXIS Bait (Plasmid #52328) and AVEXIS Prey (available at www.addgene.org), and pSecTAG2A expression vector (Thermo Fisher).

DH5α Competent Cells (Thermo Fisher) were transformed with the construct of interest and plasmids were purified using the PureLink HiPure midiprep kit (Thermo Fisher) according to manufacturer’s instructions. Plasmid constructs were transfected into Expi293F cells using the Expifectamine transfection kit (Thermo Fisher Scientific) according to manufacturer’s instructions. In brief, 3x10^6^ cells/ml at >95% viability were prepared according to culture volume required. Plasmid DNA (1 μg of plasmid DNA per mL of culture volume to transfect) was diluted in OptiMEM-I reduced serum medium and incubated with ExpiFectamine 293 Reagent (Thermo Fisher) for 20 min at room temperature (RT) and was added to Expi293F cells. Where biotinylated proteins were required, these were co-transfected additionally with biotin (100 µM) and BirA at a 1:10 ratio of plasmid DNA. After 18 hours of culture post-transfection, ExpiFectamine293 Enhancer 1 and 2 were added. Cell supernatants containing secreted protein were collected 5 days after transfection and protein expression was confirmed by western blotting.

### 2.4. Protein purification

Protein constructs containing 6HIS tags were purified from supernatants by nickel affinity chromatography using HiTrap chelating HP columns (GE Healthcare), eluting bound proteins using an imidazole gradient. Fractions containing pure expressed protein were pooled, dialysed into PBS, and sterile filtered. Protein concentration was determined by absorbance at 280 nm, calculated by each protein’s extinction coefficient.

### 2.5. Avidity-based Extracellular Interaction Screening (AVEXIS)

The Avidity-based Extracellular Interaction Screening (AVEXIS) assay is described in detail here (Kerr and Wright, 2012). Two protein libraries were generated: CCP domain-containing proteins (parasite bait) and immune targets (prey), an overview of the immune targets can be found in **Supplementary Table 1**. Streptavidin (Biolegend; 1 μg/mL) was coated on Maxisorp Nunc-Immuno 96 well plate (ThermoFisher) in carbonate buffer (0.1 M Na_2_CO_3_, 0.1 M NaHCO_3_, pH 9.6) overnight at 4°C. Plates were washed 3 times with ELISA wash buffer (PBS with 0.05% Tween 20, PBST) and blocked in block buffer 2% bovine serum albumin (BSA) in PBST for 30 min at RT. Biotinylated parasite bait proteins diluted 1:100 in block buffer were added and incubated overnight at 4°C, followed by washing. Immune prey protein was diluted to pre-determined level in block buffer and incubated overnight at 4°C. After washing, anti-beta lactamase antibody (Sigma; 1:5000 in 2% BSA PBST) was added for 1 hr at RT, followed by goat anti-rabbit IgG, HRP conjugate (Promega; 1:5000 in 2% BSA PBST) for 1 hr at RT. Plates were washed 5 times, including a 10 min final soak, then rinsed twice in water. TMB substrate was added and incubated for 20 min in the dark at RT. The reaction was stopped using 1M H_2_SO_4_, and absorbance was measured at 450 nm using a CLARIOstar Plus microplate reader (BMG Labtech).

### 2.6. ELISA binding of HpBoRB and RELM**β**

HpBoRB or RELMβ (1 µg/mL) were coated onto a 96-well plate using carbonate buffer. Plates were washed with ELISA wash buffer and blocked with block buffer for 1 hr at RT. Proteins assessed for binding include: HpBoRB, HpApyMut2, RELMβ and CD4 tag were diluted at the indicated concentration in 2% BSA PBST and incubated for 1 hr at RT. Binding was detected by probing for tags found on the respective proteins, anti-FLAG (Biolegend; 1:2500 in 2% BSA PBST), and anti-rat CD4d3+4 antibody (Bio-rad; 1:1000 in 2% BSA PBST) for 1 hr at RT. After washing, goat anti-rat IgG, HRP conjugate (Abcam; 1:5000 in 2% BSA PBST) and goat anti-mouse IgG, HRP conjugate (Bio-rad; 1:2500 in 2% BSA PBST) was added and incubated for 1 hr at RT. TMB substrate was added and incubated for 5 min in the dark at RT and the reaction was stopped using 1M H_2_SO_4._ Absorbance was measured at 450 nm and 570 nm using a CLARIOstar Plus microplate reader (BMG Labtech).

### 2.7. Competition binding assay

Streptavidin (Biolegend; 1 μg/mL) was coated on a 96-well plate using carbonate buffer and incubated overnight at 4 °C. Plate was blocked and biotinylated HpBoRB bait (1:100 in 2% BSA PBST) was added. Competing proteins HpBoRB or HpBARI, diluted to the indicated concentration in 2% BSA PBST, were co-incubated with RELMβ for 30 min at RT, then added to the plate and incubated for 1 hr at RT. RELMβ binding to immobilised HpBoRB was detected using anti-beta lactamase antibody (Sigma; 1:5000 in 2% BSA PBST), followed by goat anti-rabbit IgG, HRP conjugate (Promega; 1:5000 in 2% BSA PBST) as described previously.

### 2.8. Size exclusion chromatography

All recombinant proteins were mixed in PBS and allowed to bind for 30 min at 4 °C and then applied to a Superdex S200 10/300 GL column (Cytiva) pre-equilibrated with buffer containing 20 mM Tris-Cl pH 8.0 and 100 mM NaCl. Eluted fractions were heated at 90 °C for 5 min, then run on a 12% Tris-glycine gel under reducing conditions and transferred to nitrocellulose membranes for western blotting. Images were acquired using Chemi and IR 700 channels on a Licor Odyssey Fc.

### 2.9. Protein preparation and SPR

RELMβ was expressed as a fusion with rat CD4d3+4 followed by a BirA recognition site (for site-specific biotinylation) and a 6X-histidine tag for protein purification. NTA-purified RELMβ was further purified via size exclusion chromatography on a S200 10/300 increase (Cytiva) column equilibrated with 1X PBS to obtain aggregate-free protein. Following this procedure, 0.4-0.5 mg protein was obtained from 100 ml of Expi293F cell supernatant. The C-terminal 6X-His tagged HpBoRB protein was purified using the same protocol to yield ∼1.5-2.0 mg of protein from 100 ml of Expi293F cell supernatant. The purified RELMβ fusion protein was monobiotinylated enzymatically using BirA500 biotinylation kit (Avidity LLC) and the excess biotin was removed via PD-5 desalting column.

Experiments were performed at 25°C on a Biacore T200 instrument (GE Healthcare) using the Biotin CAPture kit (cytiva), in 1X PBS, 0.05% Tween-20. 300-400 RU RELMβ:CD4 protein was used for the binding analysis and kinetics. A twofold dilution series of HpBoRB (50 nM to 0.78 nM) was passed over the chip surface at a flow rate of 30 μl/min. Association was measured for 180 s, followed by 240 s dissociation phase. The data was processed using BIA evaluation software version 1.0 (BIAcore, GE Healthcare). Response curves were double referenced by subtracting the signal from the reference cell and averaged blank injection.

### 2.10. Statistical analysis

Data were analysed using Graphpad Prism V10.2.3. When comparing independent groups, one-way analysis of variance (ANOVA) with Dunnett’s post test was carried out. ∗∗∗∗ = P < 0.0001, ∗∗∗ = P < 0.001, ∗∗ = P < 0.01, ∗ = P < 0.05, ns = not significant (P > 0.05).

## 3. Results

### 3.1. Cataloguing of helminth CCP domain-containing proteins

Using WormBase Parasite, we collated the genomic data from 158 helminth species, and used the BioMart function to identify protein-coding genes which contained at least 1 CCP superfamily domain (Interpro IPR035976) and a signal peptide, indicating these could be secreted CCP domain proteins. Hpb contains 69 proteins with these characteristics, by far the largest number of any helminth species assessed (**Figure 1**). The next highest was the close relative of Hpb, *Nippostrongylus brasiliensis*, in which 23 signal peptide + CCP domain-containing proteins could be identified. Even considering the numbers of known CCP domain-containing immunomodulators (3 HpARIs, 3 HpBARIs, 10 TGMs = 16 total) Hpb contains a large number of extra signal peptide + CCP domain-containing proteins. Therefore, we decided to screen these Hpb proteins for further host-parasite interactions.

**Figure 1.**
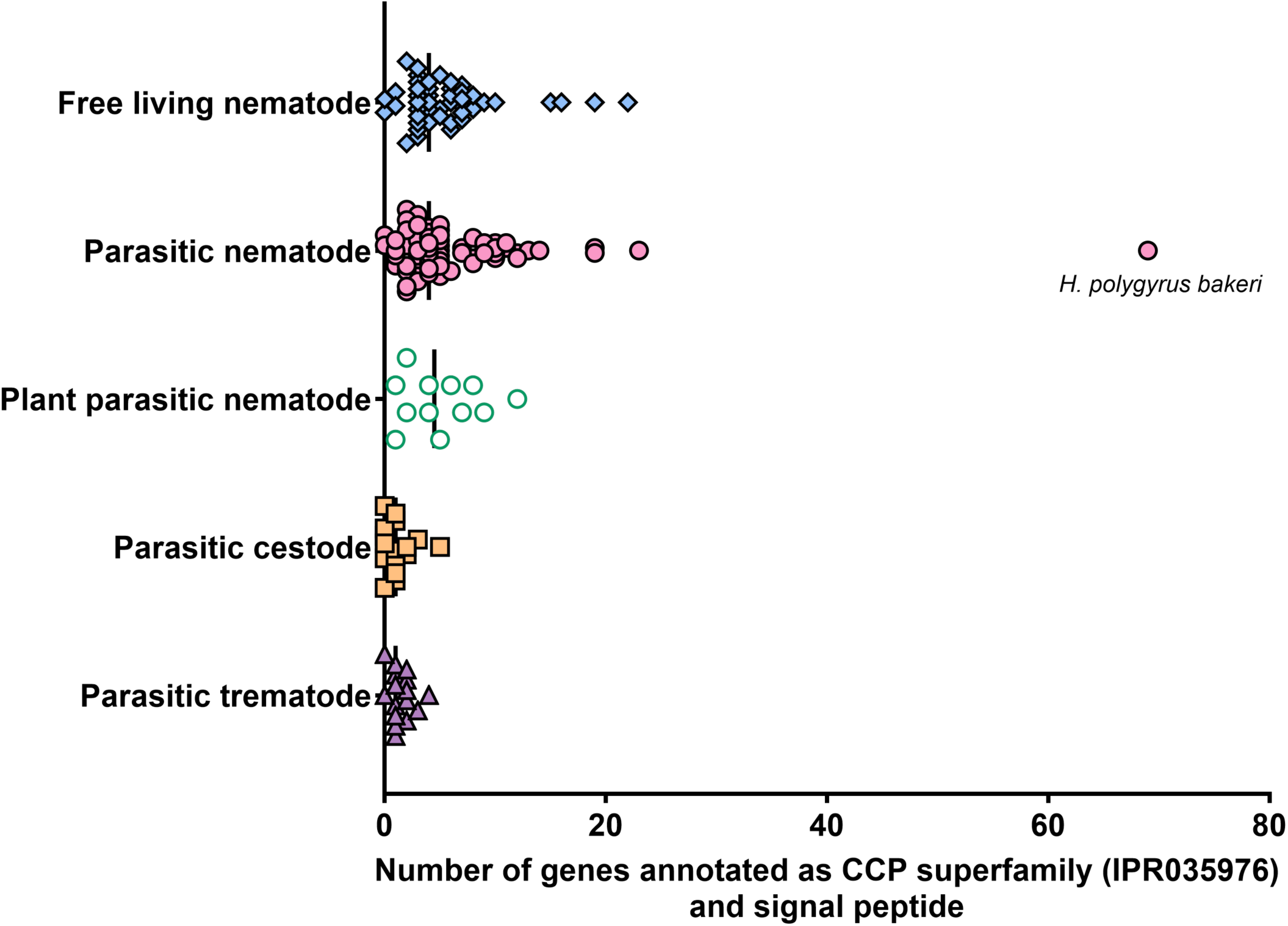
Distribution of CCP superfamily genes with a signal peptide across helminth species. Genomic data from 158 helminth species, obtained from WormBase ParaSite, were analysed to identify CCP superfamily (IPR035976) genes with a secretory signal peptide. Free living nematode (n=47), Parasitic nematode (n=67), Plant parasitic nematode (n=12), Parasitic cestode (n=16), Parasitic trematode (n=16). The line indicates the median for each group. Grubbs test was performed to detect outliers in the dataset and *H. polygyrus bakeri* was identified as an outlier.

To collate a complete list of Hpb signal peptide plus CCP domain-containing proteins in Hpb, we searched the Hpb Wormbase Parasite genomes (Bioprojects PRJEB1203 and PRJEB15396) as well as a transcriptome produced in earlier studies (Hewitson et al., 2011), and added these to the list of known immunomodulatory CCP domain-containing proteins (the HpARIs, HpBARIs and TGMs) to give a total of 112 candidates (excluding exact duplicates between the in-house transcriptome and published genomes). These translated protein sequences were aligned using Clustal Omega, and a neighbour-joining phylogenetic tree was prepared (**Figure 2**). The HpARI, HpBARI and TGM families could be clearly distinguished as forming conserved sub-families. The other CCP domain-containing proteins in the wider family contained between 1 and 16 CCP domains per protein.

**Figure 2.**
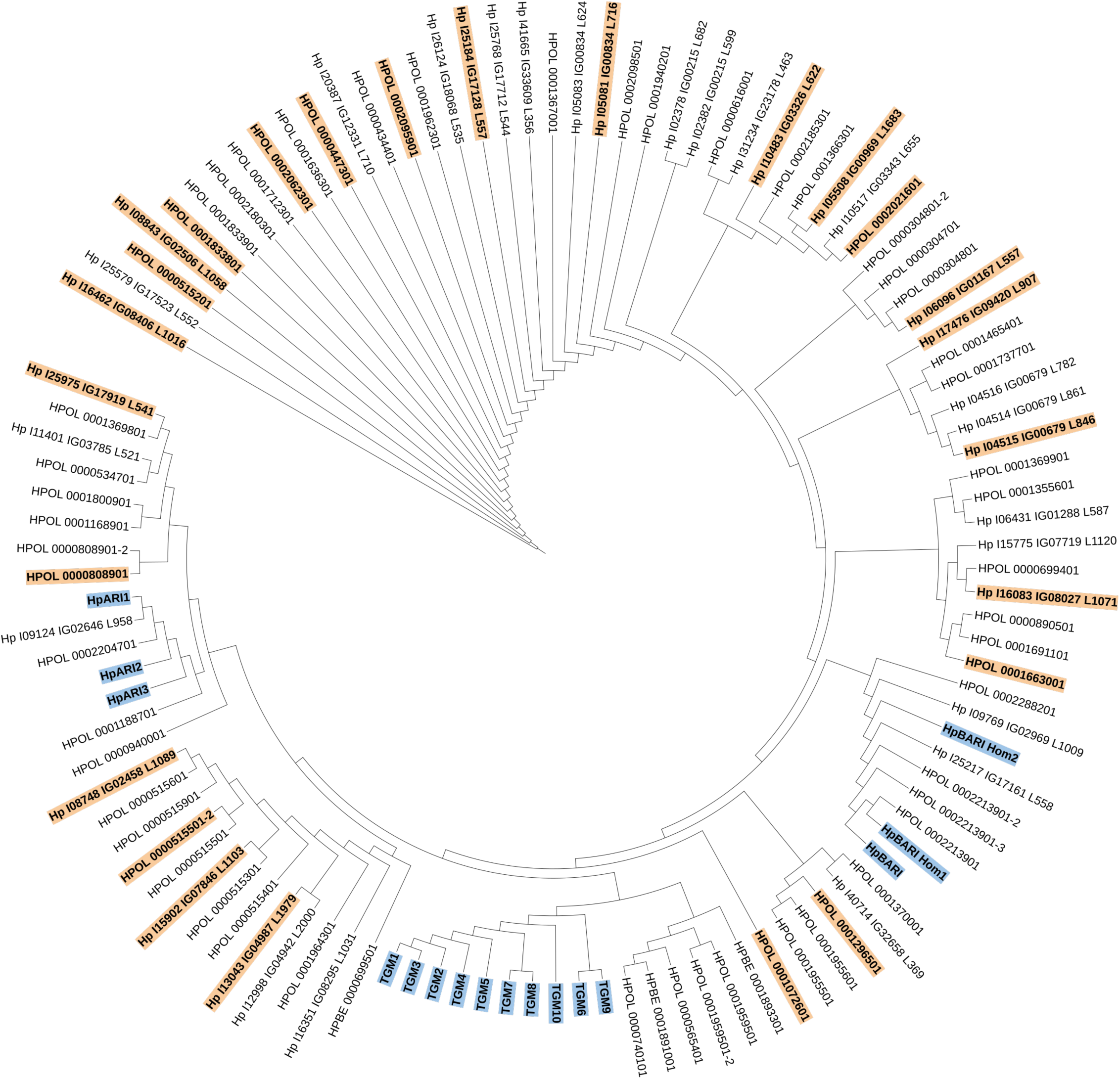
Phylogenetic tree of the CCP Superfamily proteins predicted from *H. polygyrus bakeri*. 112 CCP superfamily protein sequences were aligned using Clustal O alignment. A phylogenetic tree was constructed using the neighbour joining method and BLOSUM 62 scoring matrix based on the sequence alignment from Clustal O. Known CCP domain-containing immunomodulatory proteins are highlighted in blue. Candidates selected for expression and further testing are highlighted in orange.

### 3.2. Screening for interactions with host immune proteins

To screen proteins for potential immunomodulatory activity, we used the Avidity-based Extracellular Interaction Screening (AVEXIS) assay. This assay allows screening of biotinylated “bait” proteins (captured on a streptavidin-coated ELISA plate) against pentamerised “prey” proteins.

We selected 25 uncharacterised CCP domain-containing proteins for further investigation (**Figure 2**), along with the known immunomodulatory proteins HpBARI (Vacca et al., 2020) and TGM1 (Johnston et al., 2017). These were expressed in Expi293F cells in the bait vector as a monobiotinylated protein. These parasite baits were screened for binding to a selection of 41 host proteins associated with a type 2 immune response (**Supplementary Table 1**), including cytokines and cytokine receptor ectodomains, which were expressed in a pentamerised prey vector. Screening of parasite baits and immune preys identified the known interactions between HpBARI and ST2 (Vacca et al., 2020), and TGM1 and TGFβR2 (Johnston et al., 2017), as expected (**Figure 3A**). Interactions were also detected between 2 uncharacterised CCP domain-containing proteins: the 2 CCP domain protein HPOL_0001072601 bound to RELMβ, while the 3 CCP domain protein Hp_I7476_IG09420_L907 bound to IL1RAcP and IL-7Rα. These assays were repeated, confirming these interactions (**Figure 3B-E**). For comparison, another CCP domain-containing protein (HPOL_0000515501-2) is shown which did not interact with any immune target (**Figure 3F**). All further data from the AVEXIS assay can be found in **Supplementary Figure 1**.

**Figure 3.**
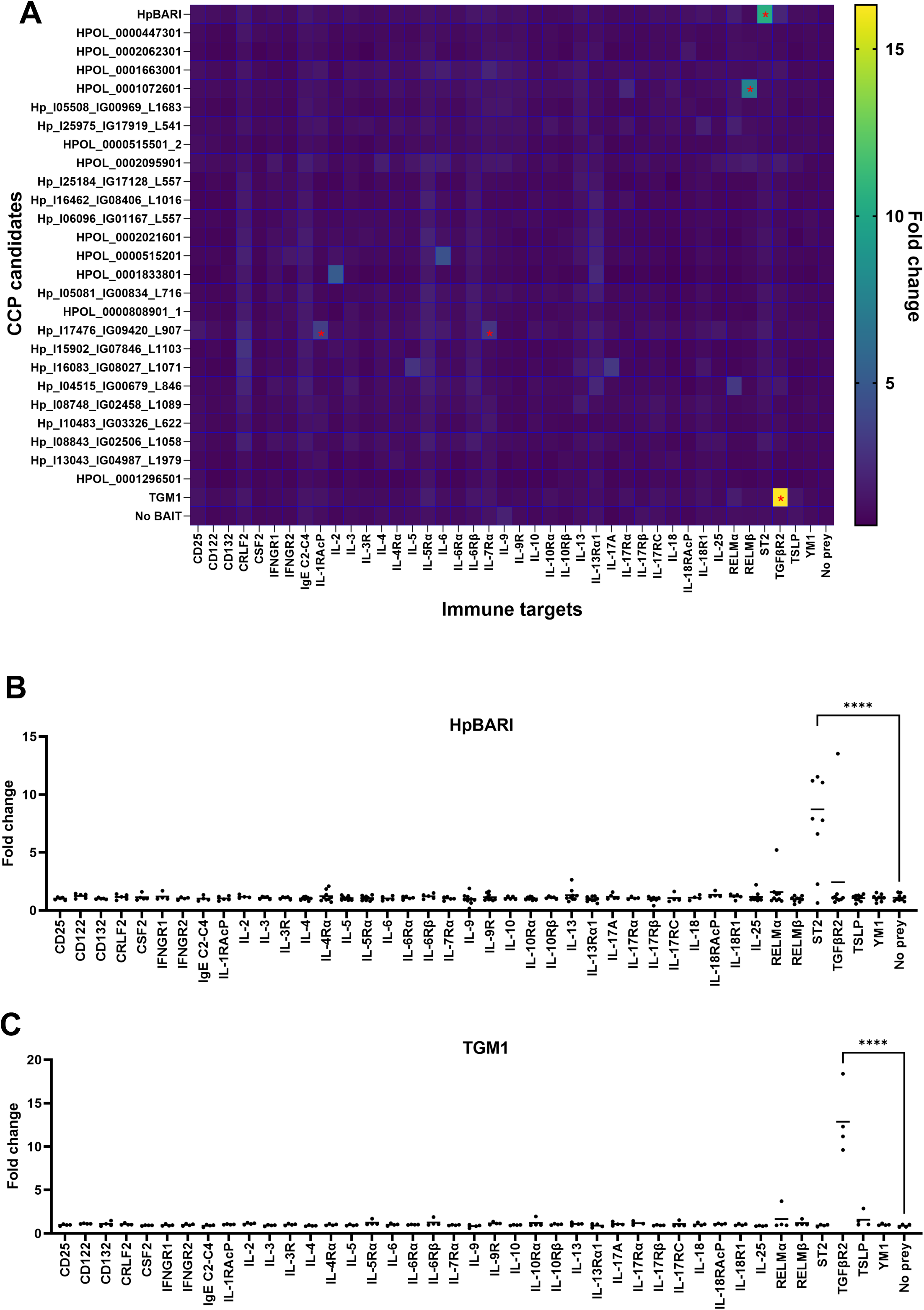
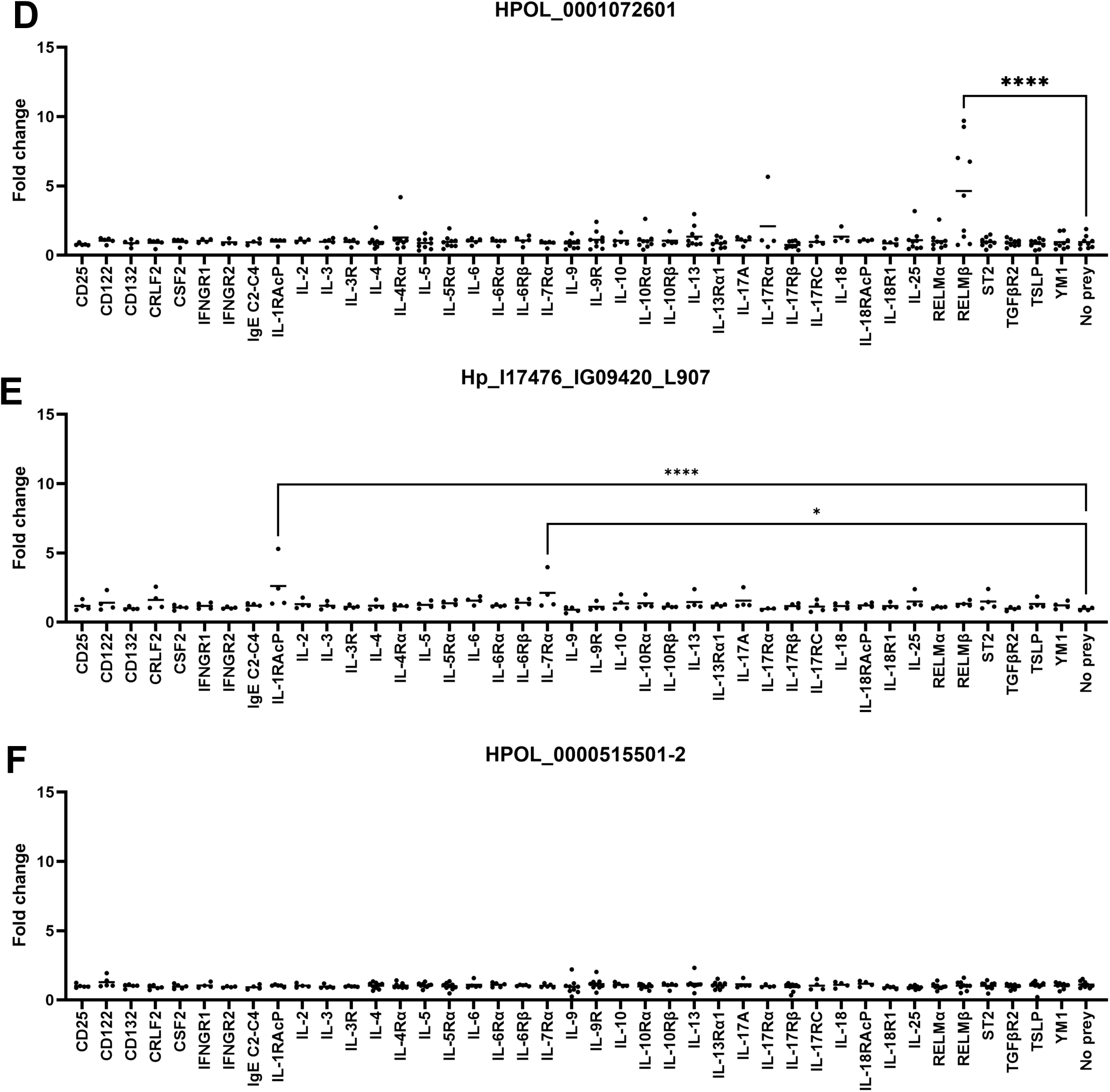
AVEXIS assay screening of CCP molecules for immune target identification. (A) AVEXIS screen heatmap showing the mean fold change for each CCP candidate against each immune prey interaction, based on 2-9 independent AVEXIS screens. Asterisks indicate statistically significant interactions. (B – F) Shows individual graphs of a CCP domain-containing protein bait screened against 41 immune preys from 2-9 independent AVEXIS screens. The Y-axis fold change was calculated by dividing the optical density for each parasite protein bait-immune prey interaction by that of the negative control no bait-immune prey interaction. The line represents the mean with each dot representing a single experiment. Results were analysed by one-way ANOVA, if a statistically significant difference was reported, a post hoc Dunnett’s multiple comparisons test to negative control was carried out. ∗ = P < 0.05, ****= p<0.0001.

As Hp_I17476_IG09420_L907 weakly bound to two unrelated surface cytokine receptors (the IL-1 family receptor IL1RAcP and the type I cytokine receptor IL-7Rα), both of which bind to multiple ligands, we decided to focus on the interaction between HPOL_0001072601 and RELMβ. HPOL_0001072601 was renamed as Binder of RELMβ (HpBoRB).

### 3.3. HpBoRB sequence, structure and expression

The HpBoRB sequence contains 2 CCP domains (each with 4 cysteine residues and a tryptophan between C_III_ and C_IV_), which is consistent with an Alphafold 3 model showing the CCP domains containing characteristic β-sheets and globular structure (**Supplementary Figure 2**). A recently-published transcriptomic dataset from Hpb (Pollo et al., 2023) was used to show that HpBoRB transcription peaks in the first week of infection, while HpBoRB protein can be detected in L4, but not adult excretory/secretory products (Hewitson et al., 2013) (**Supplementary Figure 3**). These data indicate that

HpBoRB is a 2 CCP domain protein which is expressed in tissue-dwelling L4 Hpb larvae, and not later in infection.

### 3.4. Confirmation and validation of HpBoRB as a binder of RELM**β**

To confirm binding, HpBoRB and RELMβ were subcloned into new expression vectors and purified by NTA chromatography. HpBoRB was cloned into a pSecTAG2A expression vector, while RELMβ was subcloned into the AVEXIS bait vector. When RELMβ bait was coated onto an ELISA plate, it could be bound by HpBoRB, but not a control protein (HpApyMut2, an Hpb apyrase enzyme which was mutated to ablate enzymatic activity (Berkachy et al., 2021)) expressed in the same vector (**Figure 4A**). As the RELMβ AVEXIS bait construct includes a TEV cleavage site between the rat CD4d3+4 tag and the RELMβ sequence, TEV cleavage could be used to cleave off the rat CD4d3+4 tag, which could then be purified as a control protein. Unfortunately, without the N-terminal rat CD4d3+4 solubilisation tag, tag-free RELMβ protein precipitated and could not be used for further assays. When purified HpBoRB was coated onto an ELISA plate, it interacted with RELMβ bait protein, but not the cleaved CD4 tag control (**Figure 4B**). Finally, the interaction between streptavidin-coated plate-bound biotinylated HpBoRB bait and RELMβ prey could be inhibited by competition with non-biotinylated HpBoRB protein, but not a control of non-biotinylated HpBARI protein (**Figure 4C**). These experiments confirmed the specificity of binding between HpBoRB and RELMβ, with interactions which reach saturation and indicated a subnanomolar affinity.

**Figure 4.**
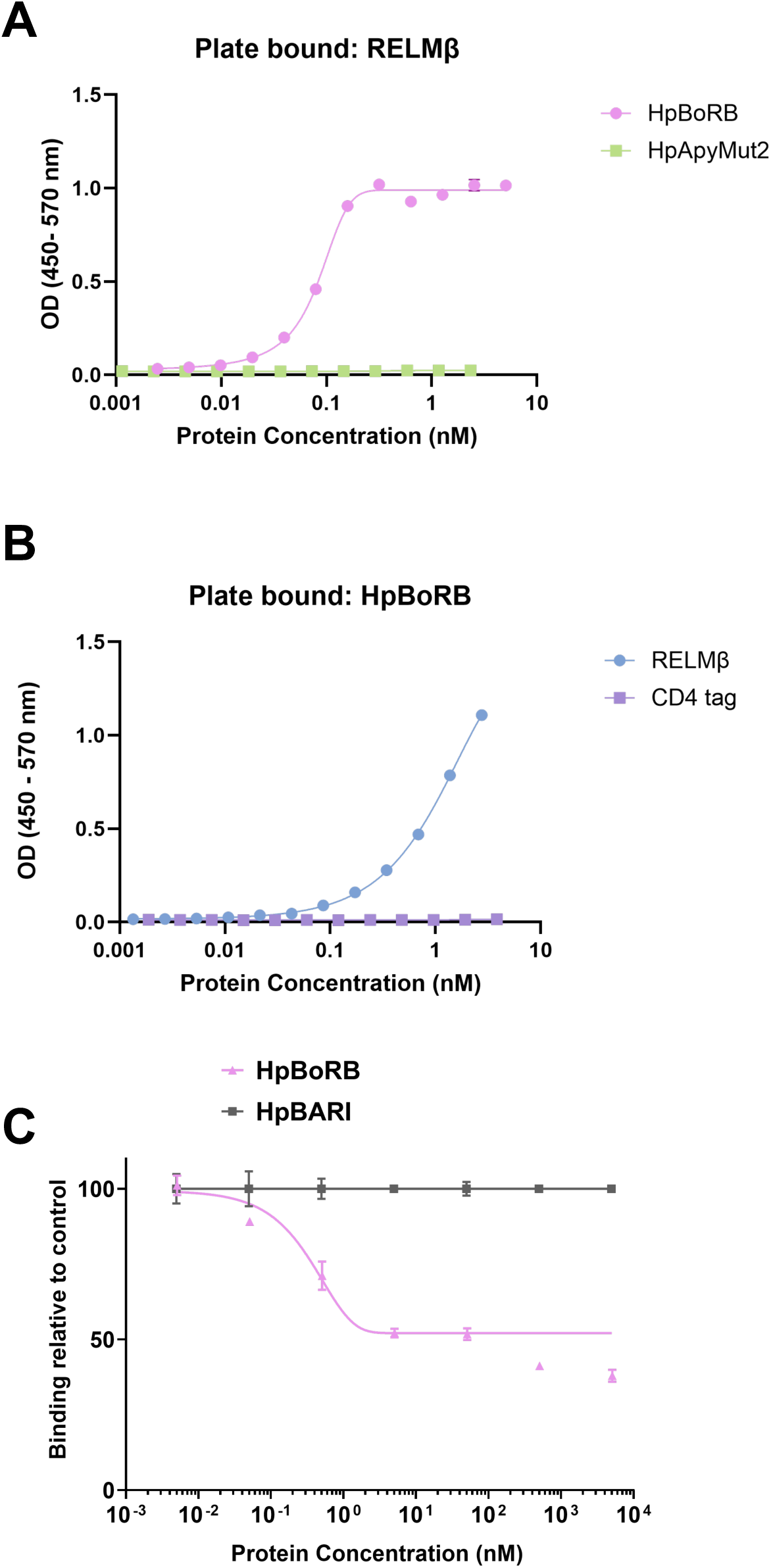
HpBoRB - RELMβ binding by ELISA. (A) RELMβ was captured on an ELISA plate and incubated with serial dilutions of HpBoRB or HpApyMut2. Protein interactions were detected using anti-FLAG antibody. SEM of 4 technical replicates shown. (B) HpBoRB was captured on an ELISA plate and incubated with serial dilutions of RELMβ or purified CD4 tag. Protein interactions were detected using anti-rat CD4 OX68 antibody. SEM of 4 technical replicates shown. (C) Competition binding of HpBoRB and RELMβ. Biotinylated HpBoRB was captured on a streptavidin-coated ELISA plate. RELMβ was co-incubated with a serial dilution of HpBoRB or HpBARI, followed by adding proteins mixtures to streptavidin-coated ELISA plates on which biotinylated HpBoRB had been captured. Protein interactions were detected using anti β-lactamase antibody. SEM of 3 technical replicates shown. All data representative of 3 independent experiments.

To further validate the interaction between HpBoRB and RELMβ in solution, HpBoRB, RELMβ, or a mixture of the two proteins were applied to a size exclusion chromatography column. A peak shift could be seen when proteins were allowed to interact, which indicated that a larger molecular mass HpBoRB-RELMβ complex was forming in solution (**Figure 5**).

**Figure 5.**
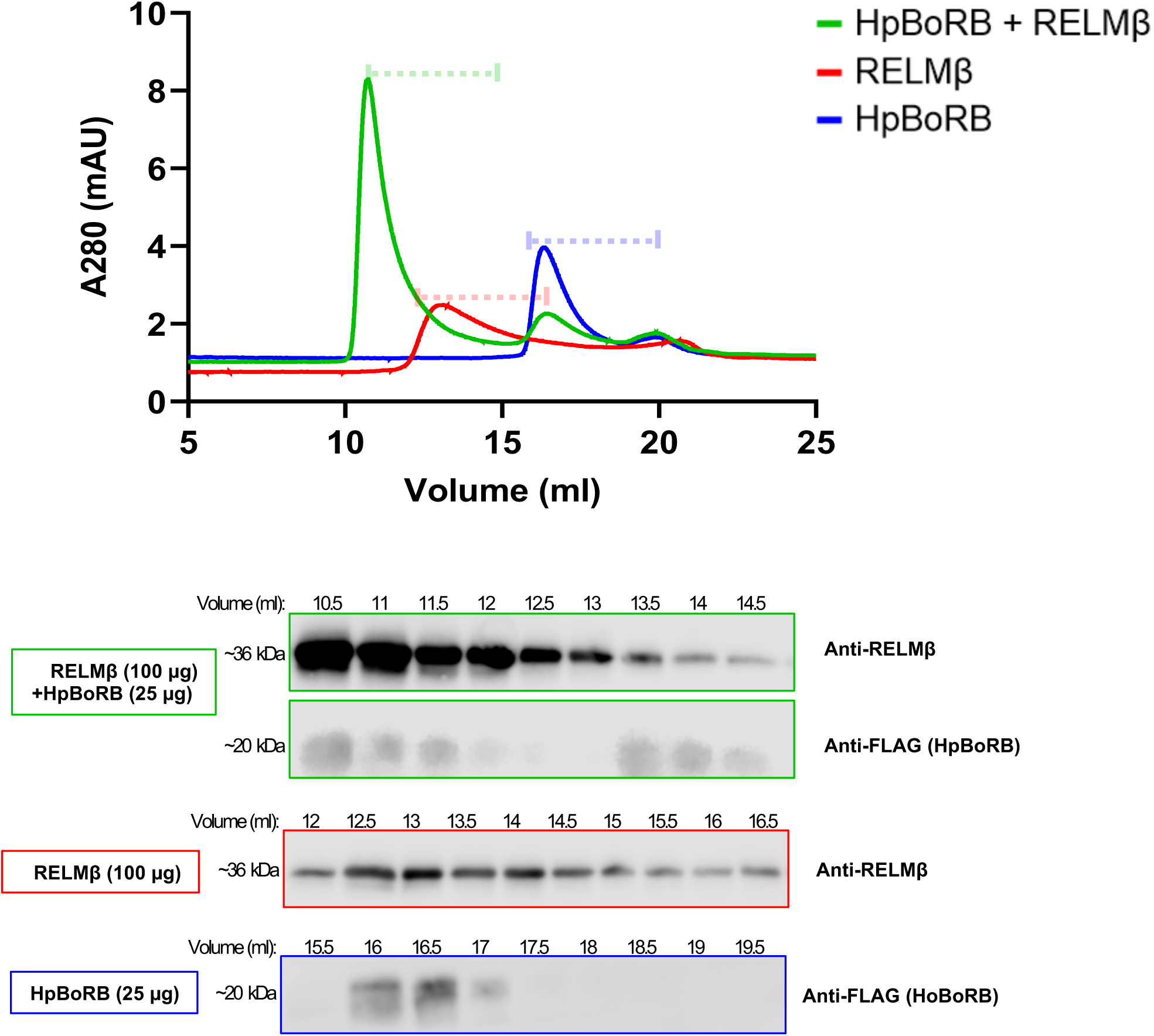
Size exclusion chromatography reveals HpBoRB binds to RELMβ. 25 μg of HpBoRB (blue), 100 μg of RELMβ (red) and 100 μg of RELMβ was added to 25 µg HpBoRB (green) and ran on a Superdex 200 Increase 10/300 GL gel filtration column. Absorbance 280 trace and 0.5 mL fractions were collected for western blot as indicated by the dotted line. Collected fractions samples probed for anti-RELMβ or anti-FLAG. In the case of sample containing HpBoRB and RELMβ, blots were probed for anti-RELMβ, stripped and re-probed for anti-FLAG. Data representative of 3 experiments.

Finally, surface plasmon resonance was used to measure the affinity of HpBoRB-RELMβ binding (**Figure 6**). Purified HpBoRB was passed over a chip coated with monobiotinylated RELMβ. HpBoRB-RELMβ binding showed a rapid on-rate and a slow off-rate, indicating a high affinity interaction with a K_D_ of <0.1 nM, and fit a 1:1 binding model. Therefore, this data confirms that HpBoRB interacts with RELMβ with high affinity.

**Figure 6.**
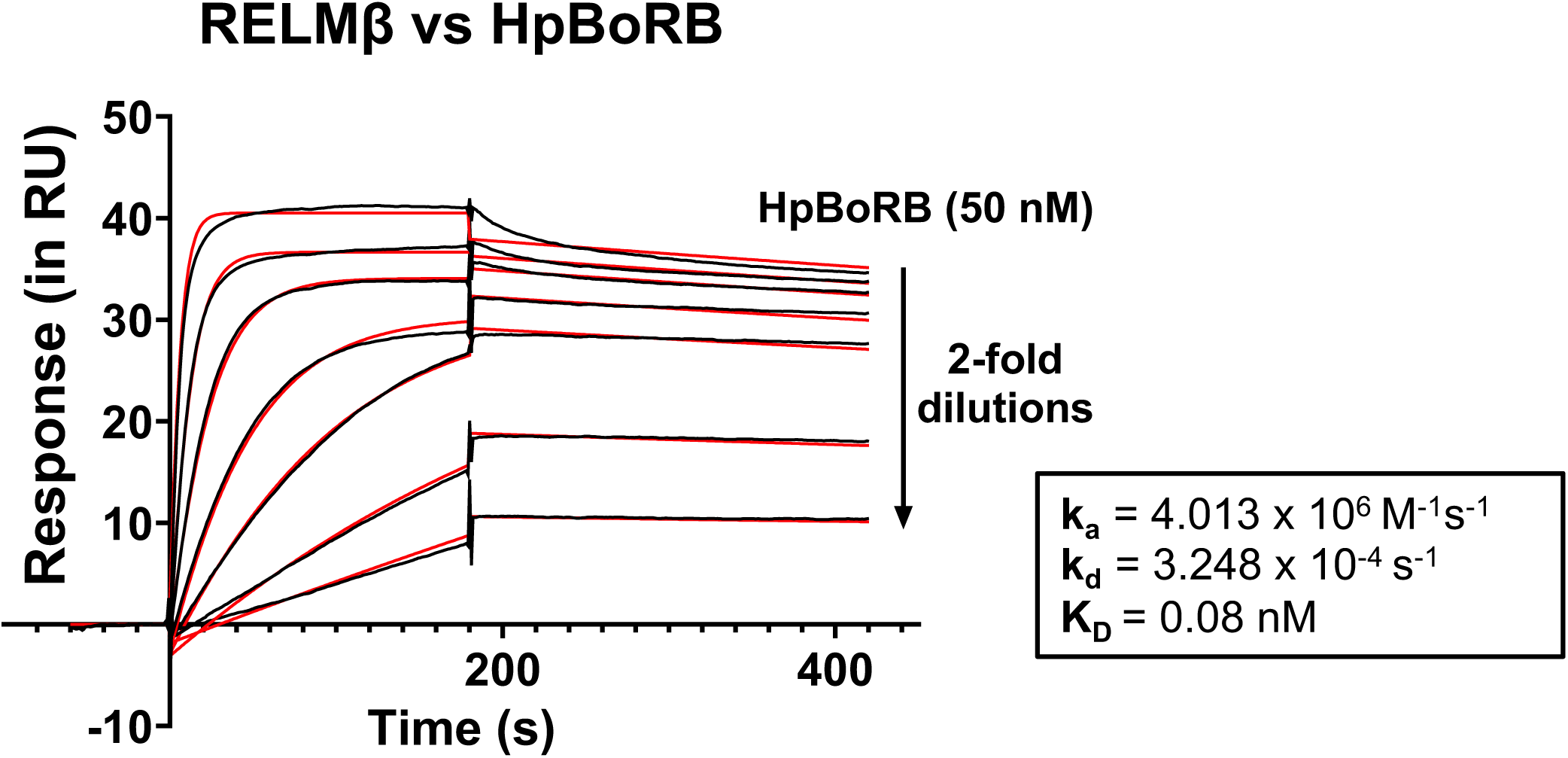
Surface plasmon resonance to determine HpBoRB - RELMβ interaction. A surface plasmon resonance sensogram showing the binding of a concentration series (two-fold dilutions from 50 nM) of HpBoRB to immobilised murine RELMβ. Black lines show SPR data and red lines fit a 1:1 binding model. Data from a single experiment.

## 4. Discussion

Parasitic helminths have an armoury of secreted proteins which they use to modulate the host immune system. Apparently uniquely, Hpb has expanded the CCP domain-containing family of proteins to act on a range of immune pathways: we show that the CCP domain-containing family is expanded in Hpb, but not in any other parasitic or free-living helminth yet characterised. This CCP domain-containing family contains the HpARI, HpBARI and TGM protein families (each with their own immunomodulatory activities), and we now show that it also contains a novel CCP domain-containing protein, HpBoRB, which binds to host RELMβ with high affinity.

Although this study did not assess the functional consequences of HpBoRB-RELMβ interactions, it is reasonable to hypothesise that HpBoRB may inhibit the activity of RELMβ. RELMβ is a member of the resistin-like molecule family of proteins (which contains resistin, RELMα, RELMβ and RELMγ), which have varied effects against bacterial and helminth infections, in metabolism, and on immune cells (Pine et al., 2018). RELMα and RELMβ are particularly strongly upregulated in type 2 immune responses, in helminth infections and asthma, however their mode of action is not well understood as no host receptor has yet been identified for either protein. However, RELMβ-deficient mice show defective clearance of several helminth infections (including Hpb), while treating adult Hpb parasites in vitro with recombinant RELMβ resulted in these worms being more rapidly ejected when transplanted to a new host (Herbert et al., 2009). Recently, RELMβ-deficient mice were found to be protected from anaphylactic reactions in food allergy models, via interactions with the intestinal microbiota and consequent reduced regulatory T cell expansion (Stephen-Victor et al., 2025). Critically, RELMβ expression was shown to reduce levels of *Lactobacillus* and *Alistipes* commensal bacteria, both of which produce indole metabolites and consequently induce regulatory T cell expansion. Thus, in RELMβ-deficient mice, or with anti-RELMβ antibody administration, *Lactobacillus* and *Alistipes* levels increase, regulatory T cells expand, and allergic immune responses are suppressed. Consistent with these observations, and our hypothesis that HpBoRB inhibits RELMβ, Hpb infection increases the abundance of *Lactobacilli* during infection, which is associated with increased regulatory T cell expansion (Reynolds et al., 2014). Therefore, through blocking both direct and indirect effects on the parasite of RELMβ, HpBoRB could aid in Hpb survival.

HpBoRB is expressed at highest levels in the tissue-dwelling larvae stage of infection (prior to day 10 of infection), similarly to the peak expression and activity of HpARI and HpBARI family members (Pollo et al., 2023; Smyth et al., 2025). Thus, this phase of infection may represent a peak of immunomodulatory activity of the parasite, inhibiting the later development of anti-parasitic responses. Recently, we showed that using a vaccination approach to block immunomodulation by HpARI and HpBARI in the early phase of infection resulted in much increased type 2 responses and effective immunity to the parasite (Smyth et al., 2025). It will be interesting to investigate the effects of blocking HpBoRB in this context.

The AVEXIS protein-protein interaction screen was limited to those immune proteins selected for cloning and expression. There may be many further interactions between these parasite secreted proteins and host partner proteins which were not included in this version of the screen. Furthermore, while the CCP domain-containing family contains a number of important immunomodulators, immunomodulation is not exclusive to this family. Recently, glutamate dehydrogenase (GDH) in Hpb secretions was shown to control type 2 immune responses through the induction of PGE2 (de Los Reyes Jimenez et al., 2020). In mammals, GDH is a metabolic enzyme, involved in the tricarboxylic acid cycle, however in Hpb (and other helminths) this enzyme has been co-opted as an immunoregulatory factor. Similarly, apyrases (extracellular ATPases) are secreted by many parasites, including Hpb, and are thought to degrade ATP, a potent extracellular damage associated molecular pattern (DAMP) (Berkachy et al., 2021). Finally, several intestinal nematodes, including *Nippostrongylus brasiliensis* and Hpb, secrete type II DNAses to degrade neutrophil extracellular traps (NETs) (Bouchery et al., 2020). As GDH, apyrases and secreted DNAses are not CCP domain proteins, but are instead enzymes which have been co-opted for immune modulation and evasion, these seem to be a separate class of host-pathogen interaction proteins, which would not have been detected by our approach.

In summary, Hpb secretes an extraordinarily large family of CCP domain-containing proteins, which have already been shown to contain two independent sets of immunomodulators. Testing the hypothesis that additional immunological functions have evolved among other members of this family, and by using a sophisticated protein-protein interaction screen, we identified HpBoRB, a novel Hpb protein which binds to RELMβ, a known player in the anti-parasite type 2 immune response which is required to eliminate helminth infection. Our future studies will investigate the role of HpBoRB in vivo, and extend our search within the CCP gene family of helminth species to fully understand their scope in modulation of the immune system.

## CRediT authorship contribution statement

**Vivien Shek**: Data curation, Investigation, Methodology, Visualization, Roles/Writing - original draft, Writing - review & editing. **Abhishek Jamwal**: Investigation, Methodology. **Danielle J Smyth**: Conceptualization, Investigation, Methodology, Supervision. **Tania Frangova**: Investigation, Methodology. **Alice R Savage**: Investigation, Methodology. **Sarah Kelly**: Investigation, Methodology. **Gavin J Wright**: Methodology, Writing - review & editing. **Rachel Toth**: Investigation, Methodology. **Rick M Maizels**: Conceptualization, Methodology, Resources, Writing - review & editing. **Matthew K Higgins**: Methodology, Supervision. **Alasdair C Ivens**: Investigation, Methodology, Supervision. **Hermelijn H Smits**: Conceptualization, Funding acquisition, Methodology, Project administration, Supervision, Writing - review & editing. **Henry J McSorley**: Conceptualization, Data curation, Funding acquisition, Investigation, Methodology, Supervision, Writing - review & editing.

## Acknowledgements

This work was funded by a grant from LONGFONDS | Accelerate as part of the AWWA project, and a Wellcome Investigator award (221914/Z/20/Z) to H.J.M. Graphic abstract created in BioRender. Mcsorley, H. (2025) https://BioRender.com/so8f8q8

## References

Almagro Armenteros, J.J., Tsirigos, K.D., Sonderby, C.K., Petersen, T.N., Winther, O., Brunak, S., von Heijne, G., Nielsen, H., 2019. SignalP 5.0 improves signal peptide predictions using deep neural networks. Nat Biotechnol 37, 420–423. doi: 10.1038/s41587-019-0036-z

Bancroft, A.J., Levy, C.W., Jowitt, T.A., Hayes, K.S., Thompson, S., McKenzie, E.A., Ball, M.D., Dubaissi, E., France, A.P., Bellina, B., Sharpe, C., Mironov, A., Brown, S.L., Cook, P.C., A, S.M., Thornton, D.J., Grencis, R.K., 2019. The major secreted protein of the whipworm parasite tethers to matrix and inhibits interleukin-13 function. Nat Commun 10, 2344. doi: 10.1038/s41467-019-09996-z

Berkachy, R., Smyth, D.J., Schnoeller, C., Harcus, Y., Maizels, R.M., Selkirk, M.E., Gounaris, K., 2021. Characterisation of the secreted apyrase family of Heligmosomoides polygyrus. Int J Parasitol 51, 39–48. doi: 10.1016/j.ijpara.2020.07.011

Blum, M., Andreeva, A., Florentino, L.C., Chuguransky, S.R., Grego, T., Hobbs, E., Pinto, B.L., Orr, A., Paysan-Lafosse, T., Ponamareva, I., Salazar, G.A., Bordin, N., Bork, P., Bridge, A., Colwell, L., Gough, J., Haft, D.H., Letunic, I., Llinares-Lopez, F., Marchler-Bauer, A., Meng-Papaxanthos, L., Mi, H., Natale, D.A., Orengo, C.A., Pandurangan, A.P., Piovesan, D., Rivoire, C., Sigrist, C.J.A., Thanki, N., Thibaud-Nissen, F., Thomas, P.D., Tosatto, S.C.E., Wu, C.H., Bateman, A., 2025. InterPro: the protein sequence classification resource in 2025. Nucleic Acids Res 53, D444–D456. doi: 10.1093/nar/gkae1082

Bouchery, T., Moyat, M., Sotillo, J., Silverstein, S., Volpe, B., Coakley, G., Tsourouktsoglou, T.D., Becker, L., Shah, K., Kulagin, M., Guiet, R., Camberis, M., Schmidt, A., Seitz, A., Giacomin, P., Le Gros, G., Papayannopoulos, V., Loukas, A., Harris, N.L., 2020. Hookworms Evade Host Immunity by Secreting a Deoxyribonuclease to Degrade Neutrophil Extracellular Traps. Cell Host Microbe 27, 277–289 e276. doi: 10.1016/j.chom.2020.01.011

Colomb, F., Jamwal, A., Ogunkanbi, A., Frangova, T., Savage, A.R., Kelly, S., Wright, G.J., Higgins, M.K., McSorley, H.J., 2024a. Heparan sulphate binding controls in vivo half-life of the HpARI protein family. eLife Sciences Publications, Ltd.

Colomb, F., McSorley, H.J., 2025. Protein families secreted by nematodes to modulate host immunity. Curr Opin Microbiol 84, 102582. doi: 10.1016/j.mib.2025.102582

Colomb, F., Ogunkanbi, A., Jamwal, A., Dong, B., Maizels, R.M., Finney, C.A.M., Wasmuth, J.D., Higgins, M.K., McSorley, H.J., 2024b. IL-33-binding HpARI family homologues with divergent effects in suppressing or enhancing type 2 immune responses. Infect Immun 92, e0039523. doi: 10.1128/iai.00395-23

de Los Reyes Jimenez, M., Lechner, A., Alessandrini, F., Bohnacker, S., Schindela, S., Trompette, A., Haimerl, P., Thomas, D., Henkel, F., Mourao, A., Geerlof, A., da Costa, C.P., Chaker, A.M., Brune, B., Nusing, R., Jakobsson, P.J., Nockher, W.A., Feige, M.J., Haslbeck, M., Ohnmacht, C., Marsland, B.J., Voehringer, D., Harris, N.L., Schmidt-Weber, C.B., Esser-von Bieren, J., 2020. An anti-inflammatory eicosanoid switch mediates the suppression of type-2 inflammation by helminth larval products. Sci Transl Med 12. doi: 10.1126/scitranslmed.aay0605

Everts, B., Hussaarts, L., Driessen, N.N., Meevissen, M.H., Schramm, G., van der Ham, A.J., van der Hoeven, B., Scholzen, T., Burgdorf, S., Mohrs, M., Pearce, E.J., Hokke, C.H., Haas, H., Smits, H.H., Yazdanbakhsh, M., 2012. Schistosome-derived omega-1 drives Th2 polarization by suppressing protein synthesis following internalization by the mannose receptor. J Exp Med 209, 1753–1767, S1751. doi: 10.1084/jem.20111381

Fumagalli, M., Sironi, M., Pozzoli, U., Ferrer-Admetlla, A., Pattini, L., Nielsen, R., 2011. Signatures of environmental genetic adaptation pinpoint pathogens as the main selective pressure through human evolution. PLoS Genet 7, e1002355. doi: 10.1371/journal.pgen.1002355

Herbert, D.R., Yang, J.Q., Hogan, S.P., Groschwitz, K., Khodoun, M., Munitz, A., Orekov, T., Perkins, C., Wang, Q., Brombacher, F., Urban, J.F., Jr., Rothenberg, M.E., Finkelman, F.D., 2009. Intestinal epithelial cell secretion of RELM-beta protects against gastrointestinal worm infection. J Exp Med 206, 2947–2957. doi: 10.1084/jem.20091268

Hewitson, J.P., Harcus, Y., Murray, J., van Agtmaal, M., Filbey, K.J., Grainger, J.R., Bridgett, S., Blaxter, M.L., Ashton, P.D., Ashford, D.A., Curwen, R.S., Wilson, R.A., Dowle, A.A., Maizels, R.M., 2011. Proteomic analysis of secretory products from the model gastrointestinal nematode Heligmosomoides polygyrus reveals dominance of venom allergen-like (VAL) proteins. J Proteomics 74, 1573–1594. doi: 10.1016/j.jprot.2011.06.002

Hewitson, J.P., Ivens, A.C., Harcus, Y., Filbey, K.J., McSorley, H.J., Murray, J., Bridgett, S., Ashford, D., Dowle, A.A., Maizels, R.M., 2013. Secretion of protective antigens by tissue-stage nematode larvae revealed by proteomic analysis and vaccination-induced sterile immunity. PLoS Pathog 9, e1003492. doi: 10.1371/journal.ppat.1003492

Howe, K.L., Bolt, B.J., Shafie, M., Kersey, P., Berriman, M., 2017. WormBase ParaSite - a comprehensive resource for helminth genomics. Mol Biochem Parasitol 215, 2–10. doi: 10.1016/j.molbiopara.2016.11.005

Jamwal, A., Colomb, F., McSorley, H.J., Higgins, M.K., 2024. Structural basis for IL-33 recognition and its antagonism by the helminth effector protein HpARI2. Nat Commun 15, 5226. doi: 10.1038/s41467-024-49550-0

Johnston, C.J.C., Smyth, D.J., Kodali, R.B., White, M.P.J., Harcus, Y., Filbey, K.J., Hewitson, J.P., Hinck, C.S., Ivens, A., Kemter, A.M., Kildemoes, A.O., Le Bihan, T., Soares, D.C., Anderton, S.M., Brenn, T., Wigmore, S.J., Woodcock, H.V., Chambers, R.C., Hinck, A.P., McSorley, H.J., Maizels, R.M., 2017. A structurally distinct TGF-beta mimic from an intestinal helminth parasite potently induces regulatory T cells. Nat Commun 8, 1741. doi: 10.1038/s41467-017-01886-6

Kerr, J.S., Wright, G.J., 2012. Avidity-based extracellular interaction screening (AVEXIS) for the scalable detection of low-affinity extracellular receptor-ligand interactions. J Vis Exp, e3881. doi: 10.3791/3881

Lambrecht, B.N., Hammad, H., 2017. The immunology of the allergy epidemic and the hygiene hypothesis. Nat Immunol 18, 1076–1083. doi: 10.1038/ni.3829

Maizels, R.M., McSorley, H.J., Smits, H.H., Ten Dijke, P., Hinck, A.P., 2025. Cytokines from parasites: manipulating host responses by molecular mimicry. Biochem J 482. doi: 10.1042/BCJ20253061

Maizels, R.M., Smits, H.H., McSorley, H.J., 2018. Modulation of Host Immunity by Helminths: The Expanding Repertoire of Parasite Effector Molecules. Immunity 49, 801–818. doi: 10.1016/j.immuni.2018.10.016

Navarro, S., Pickering, D.A., Ferreira, I.B., Jones, L., Ryan, S., Troy, S., Leech, A., Hotez, P.J., Zhan, B., Laha, T., Prentice, R., Sparwasser, T., Croese, J., Engwerda, C.R., Upham, J.W., Julia, V., Giacomin, P.R., Loukas, A., 2016. Hookworm recombinant protein promotes regulatory T cell responses that suppress experimental asthma. Sci Transl Med 8, 362ra143. doi: 10.1126/scitranslmed.aaf8807

Nisbet, A.J., McNeilly, T.N., Wildblood, L.A., Morrison, A.A., Bartley, D.J., Bartley, Y., Longhi, C., McKendrick, I.J., Palarea-Albaladejo, J., Matthews, J.B., 2013. Successful immunization against a parasitic nematode by vaccination with recombinant proteins. Vaccine 31, 4017–4023. doi: 10.1016/j.vaccine.2013.05.026

Osbourn, M., Soares, D.C., Vacca, F., Cohen, E.S., Scott, I.C., Gregory, W.F., Smyth, D.J., Toivakka, M., Kemter, A.M., le Bihan, T., Wear, M., Hoving, D., Filbey, K.J., Hewitson, J.P., Henderson, H., Gonzalez-Ciscar, A., Errington, C., Vermeren, S., Astier, A.L., Wallace, W.A., Schwarze, J., Ivens, A.C., Maizels, R.M., McSorley, H.J., 2017. HpARI Protein Secreted by a Helminth Parasite Suppresses Interleukin-33. Immunity 47, 739–751 e735. doi: 10.1016/j.immuni.2017.09.015

Pine, G.M., Batugedara, H.M., Nair, M.G., 2018. Here, there and everywhere: Resistin-like molecules in infection, inflammation, and metabolic disorders. Cytokine 110, 442–451. doi: 10.1016/j.cyto.2018.05.014

Pollo, S.M.J., Leon-Coria, A., Liu, H., Cruces-Gonzalez, D., Finney, C.A.M., Wasmuth, J.D., 2023. Transcriptional patterns of sexual dimorphism and in host developmental programs in the model parasitic nematode Heligmosomoides bakeri. Parasit Vectors 16, 171. doi: 10.1186/s13071-023-05785-2

Reynolds, L.A., Smith, K.A., Filbey, K.J., Harcus, Y., Hewitson, J.P., Redpath, S.A., Valdez, Y., Yebra, M.J., Finlay, B.B., Maizels, R.M., 2014. Commensal-pathogen interactions in the intestinal tract: lactobacilli promote infection with, and are promoted by, helminth parasites. Gut Microbes 5, 522–532. doi: 10.4161/gmic.32155

Ryan, S.M., Ruscher, R., Johnston, W.A., Pickering, D.A., Kennedy, M.W., Smith, B.O., Jones, L., Buitrago, G., Field, M.A., Esterman, A.J., McHugh, C.P., Browne, D.J., Cooper, M.M., Ryan, R.Y.M., Doolan, D.L., Engwerda, C.R., Miles, K., Mitreva, M., Croese, J., Rahman, T., Alexandrov, K., Giacomin, P.R., Loukas, A., 2022. Novel antiinflammatory biologics shaped by parasite-host coevolution. Proc Natl Acad Sci U S A 119, e2202795119. doi: 10.1073/pnas.2202795119

Singh, S.P., Smyth, D.J., Cunningham, K.T., Mukundan, A., Byeon, C.H., Hinck, C.S., White, M.P.J., Ciancia, C., Wasowska, N., Sanders, A., Jin, R., White, R.F., Lilla, S., Zanivan, S., Schoenherr, C., Inman, G.J., van Dinther, M., Ten Dijke, P., Hinck, A.P., Maizels, R.M., 2025. The TGF-beta mimic TGM4 achieves cell specificity through combinatorial surface co-receptor binding. EMBO Rep 26, 218–244. doi: 10.1038/s44319-024-00323-2

Smyth, D.J., Harcus, Y., White, M.P.J., Gregory, W.F., Nahler, J., Stephens, I., Toke-Bjolgerud, E., Hewitson, J.P., Ivens, A., McSorley, H.J., Maizels, R.M., 2018. TGF-beta mimic proteins form an extended gene family in the murine parasite Heligmosomoides polygyrus. Int J Parasitol 48, 379–385. doi: 10.1016/j.ijpara.2017.12.004

Smyth, D.J., Hodge, S.H., Ong, N.W.P., Richards, J., Colomb, F., Di Carmine, S., Shek, V., Frangova, T., Poveda, M.C., Maizels, R.M., McSorley, H.J., 2025. Vaccination against helminth IL-33 modulators permits immune-mediated parasite ejection. Cell Rep 44, 115721. doi: 10.1016/j.celrep.2025.115721

Stephen-Victor, E., Kuziel, G.A., Martinez-Blanco, M., Jugder, B.E., Benamar, M., Wang, Z., Chen, Q., Lozano, G.L., Abdel-Gadir, A., Cui, Y., Fong, J., Saint-Denis, E., Chang, I., Nadeau, K.C., Phipatanakul, W., Zhang, A., Farraj, F.A., Holder-Niles, F., Zeve, D., Breault, D.T., Schmitz-Abe, K., Rachid, R., Crestani, E., Rakoff-Nahoum, S., Chatila, T.A., 2025. RELMbeta sets the threshold for microbiome-dependent oral tolerance. Nature 638, 760–768. doi: 10.1038/s41586-024-08440-7

Vacca, F., Chauche, C., Jamwal, A., Hinchy, E.C., Heieis, G., Webster, H., Ogunkanbi, A., Sekne, Z., Gregory, W.F., Wear, M., Perona-Wright, G., Higgins, M.K., Nys, J.A., Cohen, E.S., McSorley, H.J., 2020. A helminth-derived suppressor of ST2 blocks allergic responses. Elife 9. doi: 10.7554/eLife.54017

van Dinther, M., Cunningham, K.T., Singh, S.P., White, M.P.J., Campion, T., Ciancia, C., van Veelen, P.A., de Ru, A.H., Gonzalez-Prieto, R., Mukundan, A., Byeon, C.H., Staggers, S.R., Hinck, C.S., Hinck, A.P., Dijke, P.T., Maizels, R.M., 2023. CD44 acts as a coreceptor for cell-specific enhancement of signaling and regulatory T cell induction by TGM1, a parasite TGF-beta mimic. Proc Natl Acad Sci U S A 120, e2302370120. doi: 10.1073/pnas.2302370120

White, M.P.J., Smyth, D.J., Cook, L., Ziegler, S.F., Levings, M.K., Maizels, R.M., 2021. The parasite cytokine mimic Hp-TGM potently replicates the regulatory effects of TGF-beta on murine CD4(+) T cells. Immunol Cell Biol 99, 848–864. doi: 10.1111/imcb.12479

White, S.E., Schwartze, T.A., Mukundan, A., Schoenherr, C., Singh, S.P., van Dinther, M., Cunningham, K.T., White, M.P.J., Campion, T., Pritchard, J., Hinck, C.S., Ten Dijke, P., Inman, G.J., Maizels, R.M., Hinck, A.P., 2025. TGM6 is a helminth secretory product that mimics TGF-beta binding to TGFBR2 to antagonize signaling in fibroblasts. Nat Commun 16, 1847. doi: 10.1038/s41467-025-56954-z

