## Supplementary figures for "HpBoRB, a helminth-derived CCP domain protein which binds RELMβ"

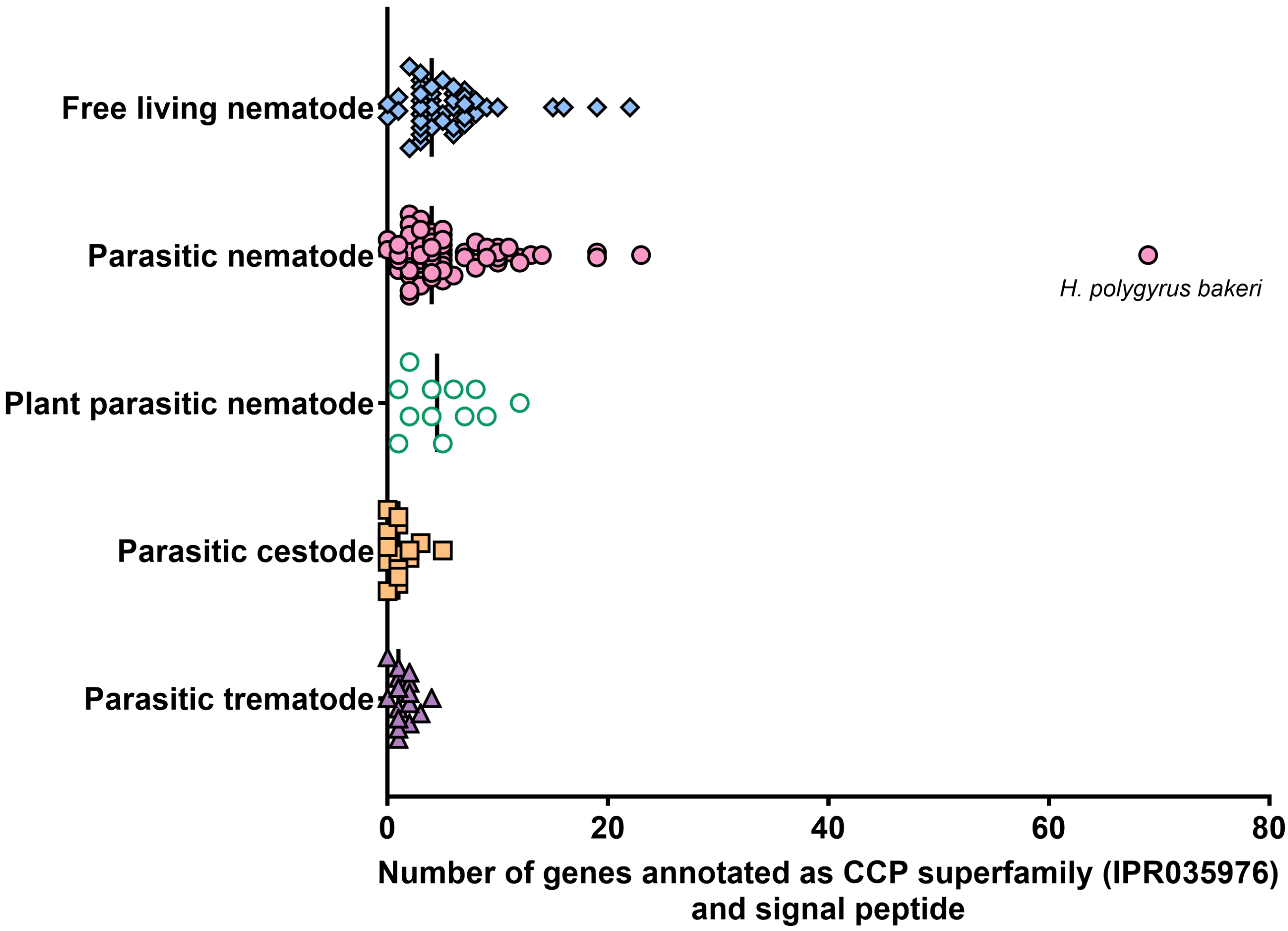


**Figure 1 - Distribution of CCP superfamily genes with a signal peptide across helminth species.** Genomic data from 158 helminth species, obtained from WormBase ParaSite, were analysed to identify CCP superfamily (IPR035976) genes with a secretory signal peptide. Free living nematode (n=47), Parasitic nematode (n=67), Plant parasitic nematode (n=12), Parasitic cestode (n=16), Parasitic trematode (n=16). The line indicates the median for each group. Grubbs test was performed to detect outliers in the dataset and *H. polygyrus bakeri* was identified as an outlier.

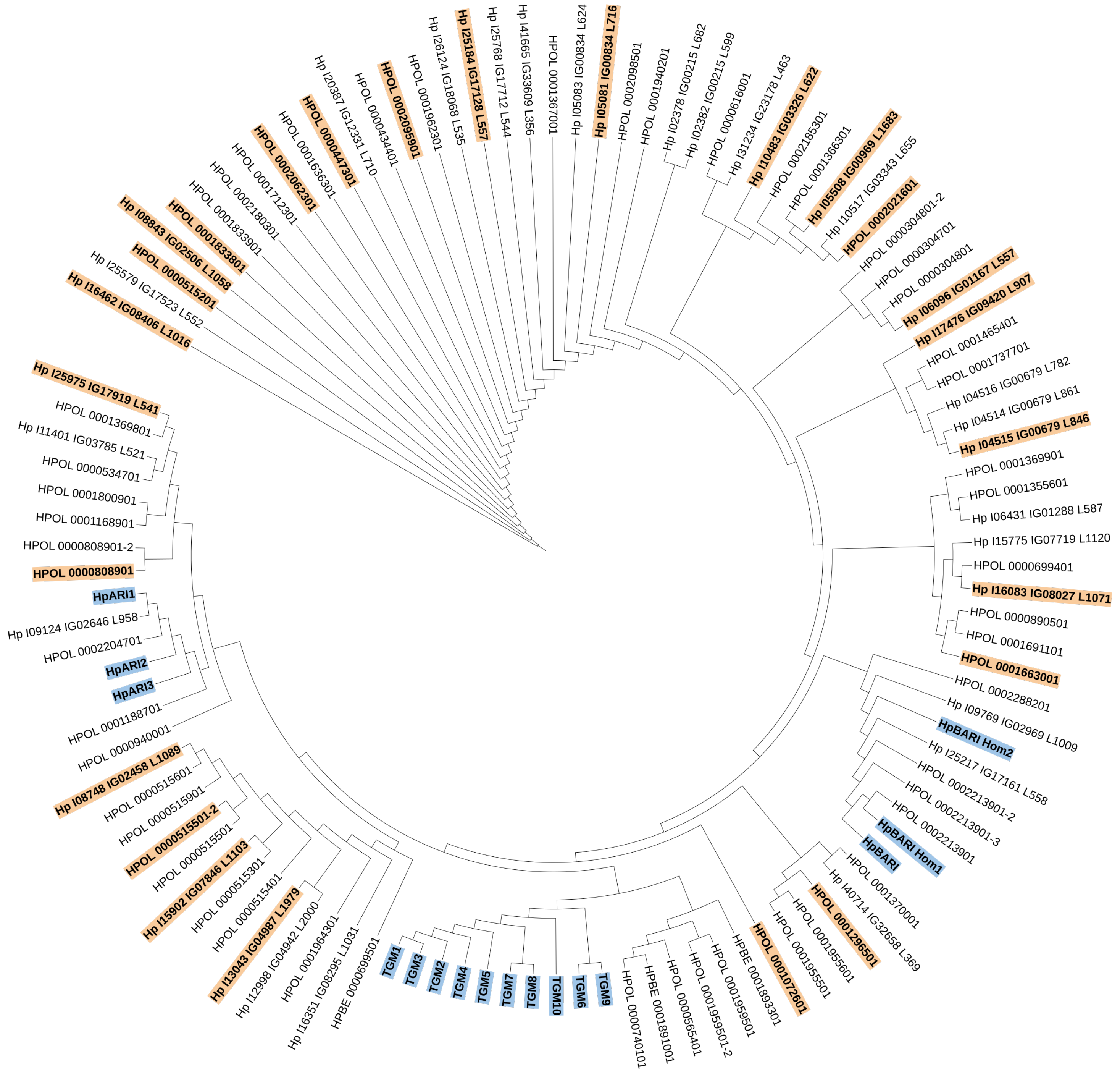


**Figure 2 – Phylogenetic tree of the CCP Superfamily proteins predicted from *H. polygyrus bakeri***

112 CCP superfamily protein sequences were aligned using Clustal O alignment. A phylogenetic tree was constructed using the neighbour joining method and BLOSUM 62 scoring matrix based on the sequence alignment from Clustal O. Known CCP domain-containing immunomodulatory proteins are highlighted in blue. Candidates selected for expression and further testing are highlighted in orange.

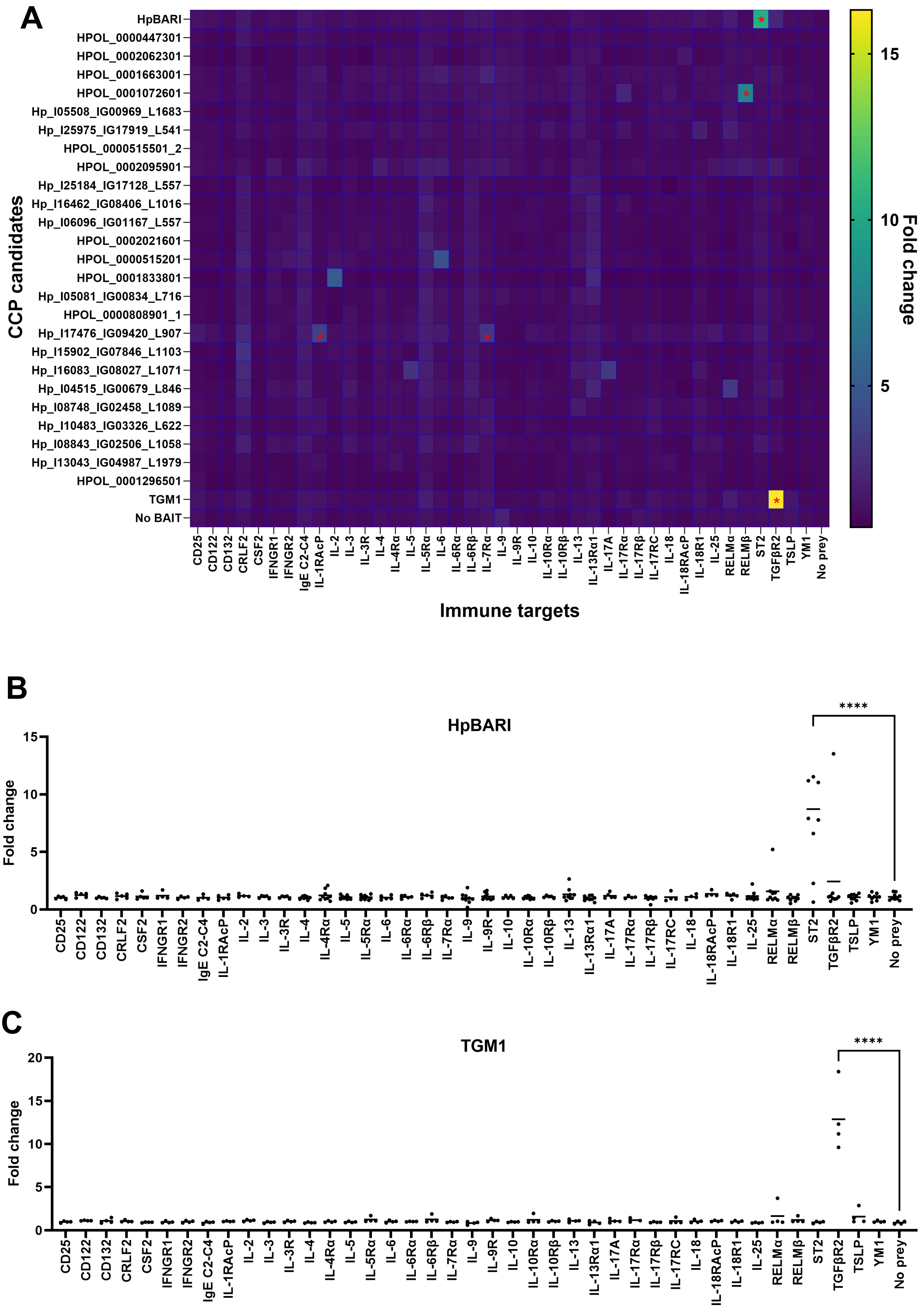
A
CCP candidates
HpBARI
HPOL_0000447301
HPOL_0002062301
HPOL_0001663001
HPOL_0001072601
Hp_I05508_IG00969_L1683
Hp_I25975_IG17919_L541
HPOL_0000515501_2
HPOL_0002095901
Hp_I25184_IG17128_L557
Hp_I16462_IG08406_L1016
Hp_I06096_IG01167_L557
HPOL_0002021601
HPOL_0000515201
HPOL_0001833801
Hp_I05081_IG00834_L716
HPOL_0000808901_1
Hp_I17476_IG09420_L907
Hp_I15902_IG07846_L1103
Hp_I16083_IG08027_L1071
Hp_I04515_IG00679_L846
Hp_I08748_IG02458_L1089
Hp_I10483_IG03326_L622
Hp_I08843_IG02506_L1058
Hp_I13043_IG04987_L1979
HPOL_0001296501
TGM1
No BAIT
CD25
CD122
CD132
CRLF2
CSF2
IFNGR1
IFNGR2
IgE C2-C4
IL-1RAcP
IL-2
IL-3
IL-3R
IL-4
IL-4Rα
IL-5
IL-5Rα
IL-6
IL-6Rα
IL-6Rβ
IL-7Rα
IL-9
IL-9R
IL-10
IL-10Rα
IL-10Rβ
IL-13
IL-13Rα1
IL-17A
IL-17Rα
IL-17Rβ
IL-17RC
IL-18
IL-18RAcP
IL-18R1
IL-25
RELMα
RELMβ
ST2
TGFβR2
TSLP
YM1
No prey
Immune targets
Fold change
15
10
5
B
HpBARI
Fold change
15
10
5
0
****
C
TGM1
Fold change
20
15
10
5
0
****

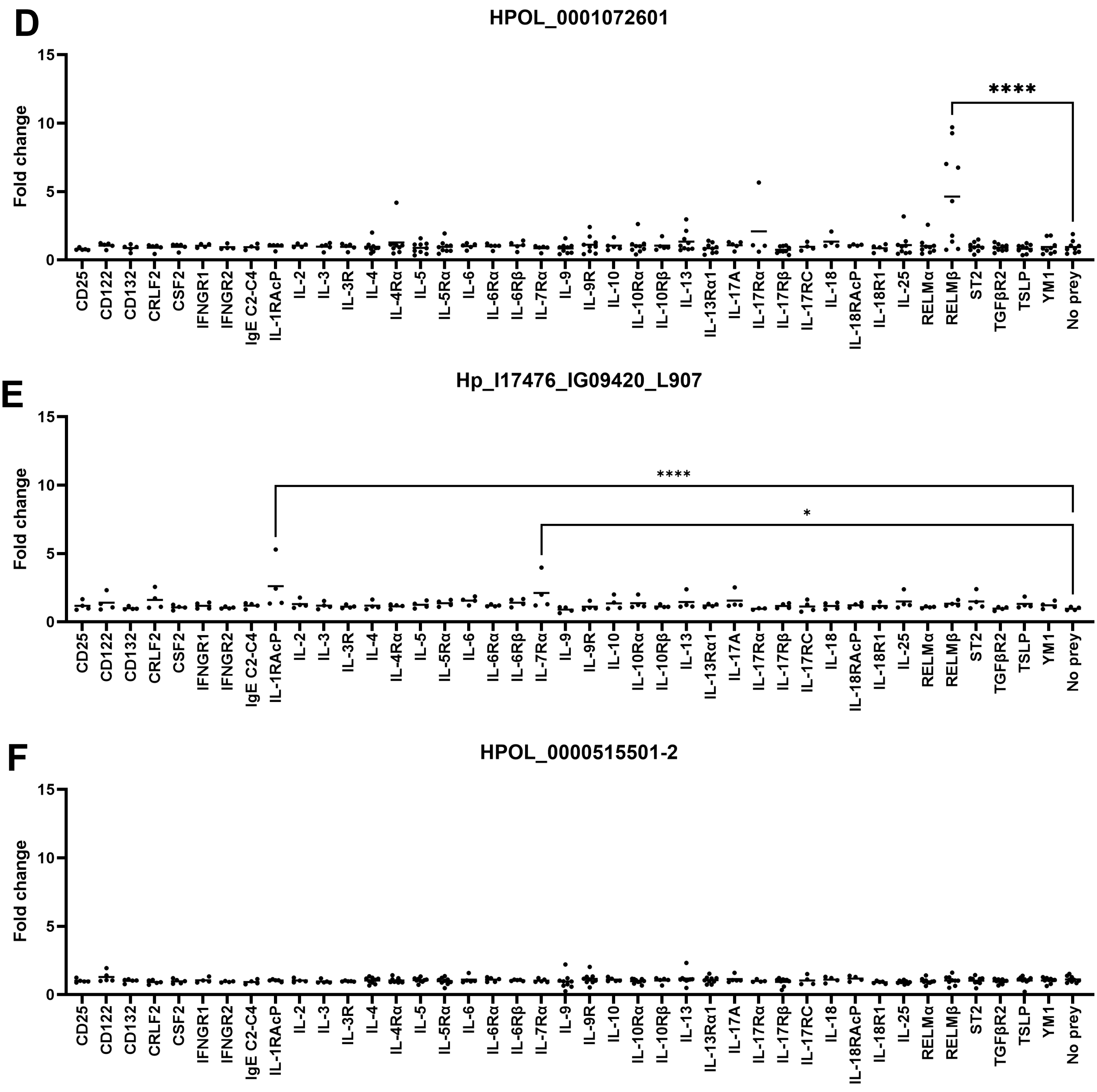


**Figure 3 - AVEXIS assay screening of CCP molecules for immune target identification.**
(A) AVEXIS screen heatmap showing the mean fold change for each CCP candidate against each immune prey interaction, based on 2-9 independent AVEXIS screens. Asterisks indicate statistically significant interactions.
(B – F) Shows individual graphs of a CCP domain-containing protein bait screened against 41 immune preys from 2-9 independent AVEXIS screens.

The Y-axis fold change was calculated by dividing the optical density for each parasite protein bait-immune prey interaction by that of the negative control no bait-immune prey interaction. The line represents the mean with each dot representing a single experiment. Results were analysed by one-way ANOVA, if a statistically significant difference was reported, a post hoc Dunnett's multiple comparisons test to negative control was carried out. ∗ = $P < 0.05$, ****= $p<0.0001$.

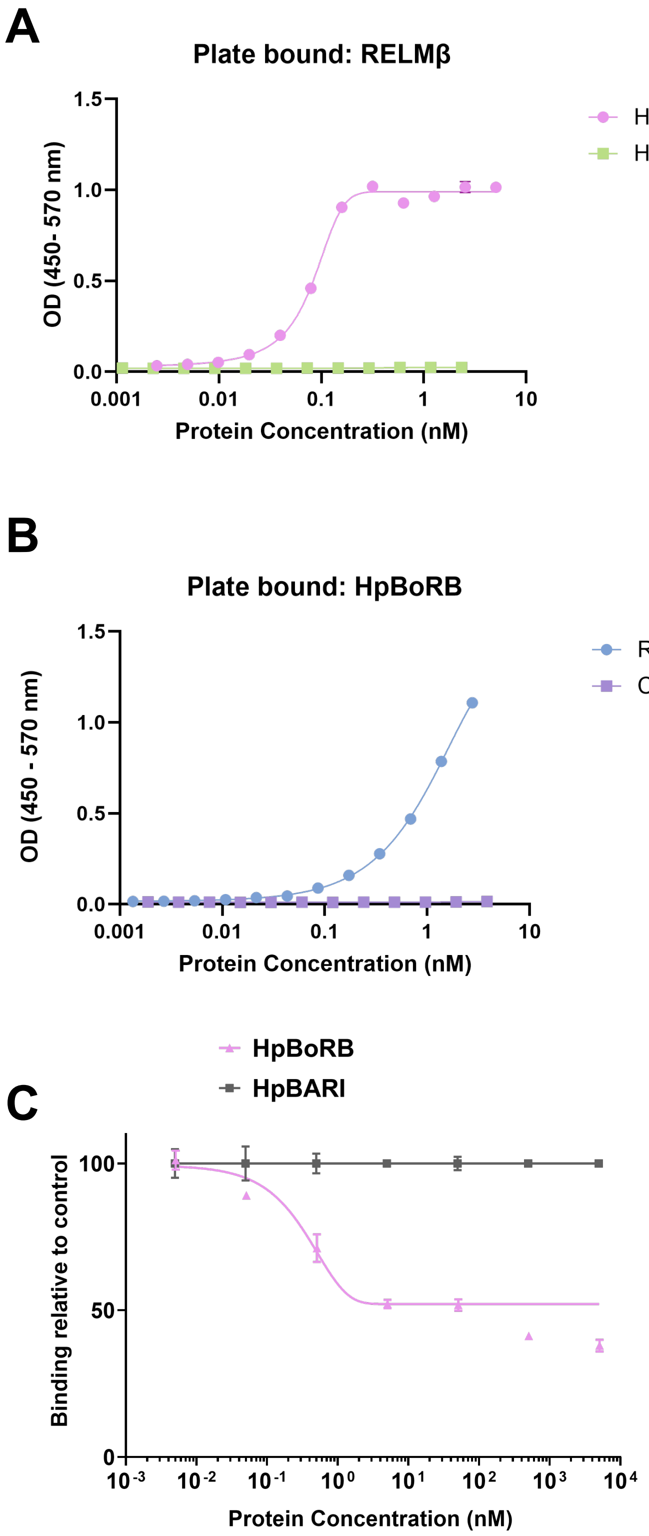


**Figure 4 HpBoRB - RELMβ binding by ELISA**

(A) RELMβ was captured on an ELISA plate and incubated with serial dilutions of HpBoRB or HpApyMut2. Protein interactions were detected using anti-FLAG antibody. SEM of 4 technical replicates shown.

(B) HpBoRB was captured on an ELISA plate and incubated with serial dilutions of RELMβ or purified CD4 tag. Protein interactions were detected using anti-rat CD4 OX68 antibody. SEM of 4 technical replicates shown.

(C) Competition binding of HpBoRB and RELMβ. Biotinylated HpBoRB was captured on a streptavidin-coated ELISA plate. RELMβ was co-incubated with a serial dilution of HpBoRB or HpBARI, followed by adding proteins mixtures to streptavidin-coated ELISA plates on which biotinylated HpBoRB had been captured. Protein interactions were detected using anti β-lactamase antibody. SEM of 3 technical replicates shown.

All data representative of 3 independent experiments.

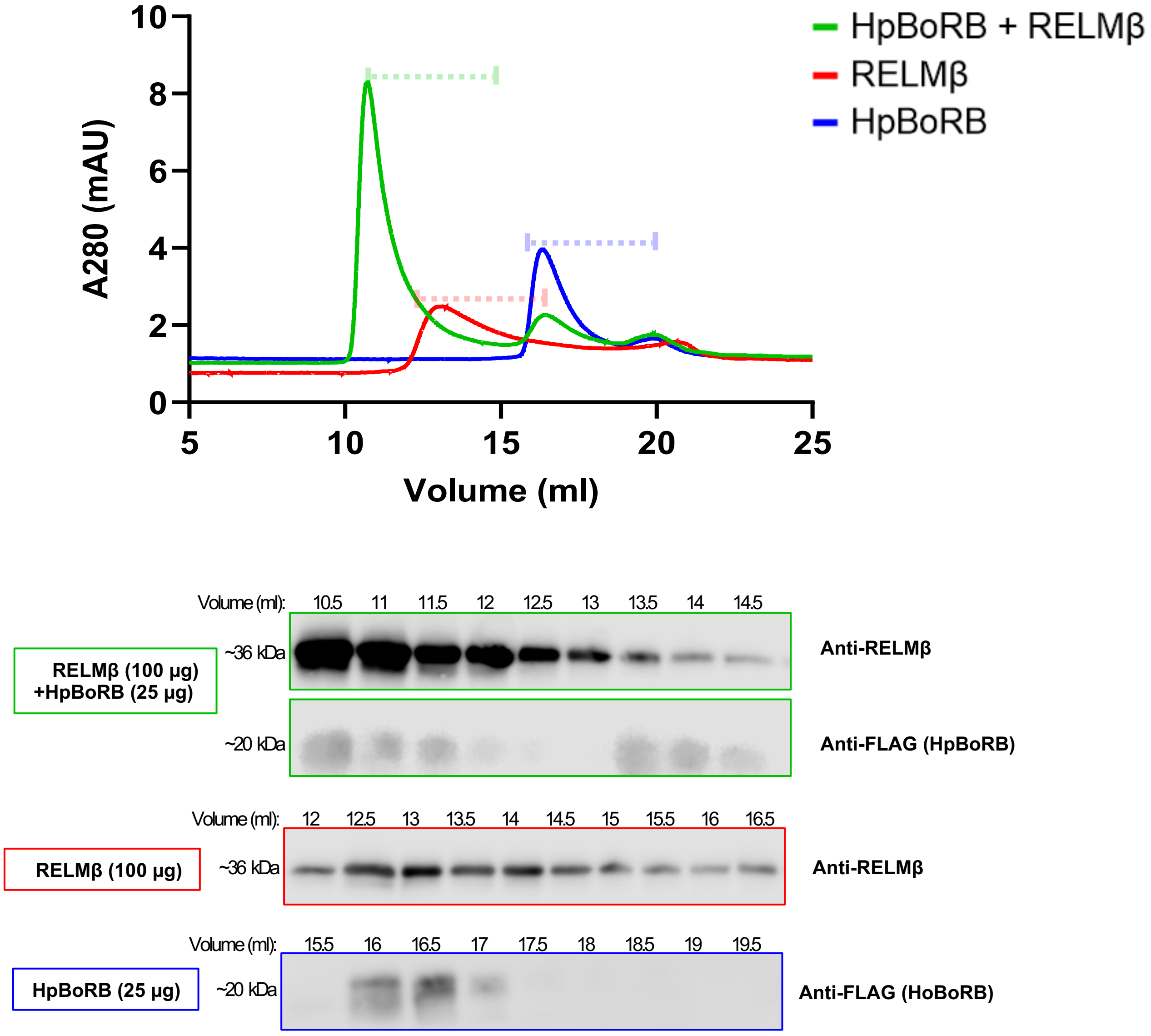


**Figure 5 - Size exclusion chromatography reveals HpBoRB binds to RELMβ**
25 µg of HpBoRB (blue), 100 µg of RELMβ (red) and 100 µg of RELMβ was added to 25 µg HpBoRB (green) and ran on a Superdex 200 Increase 10/300 GL gel filtration column. Absorbance 280 trace and 0.5 mL fractions were collected for western blot as indicated by the dotted line. Collected fractions samples probed for anti-RELMβ or anti-FLAG. In the case of sample containing HpBoRB and RELMβ, blots were probed for anti-RELMβ, stripped and re-probed for anti-FLAG. Data representative of 3 experiments.

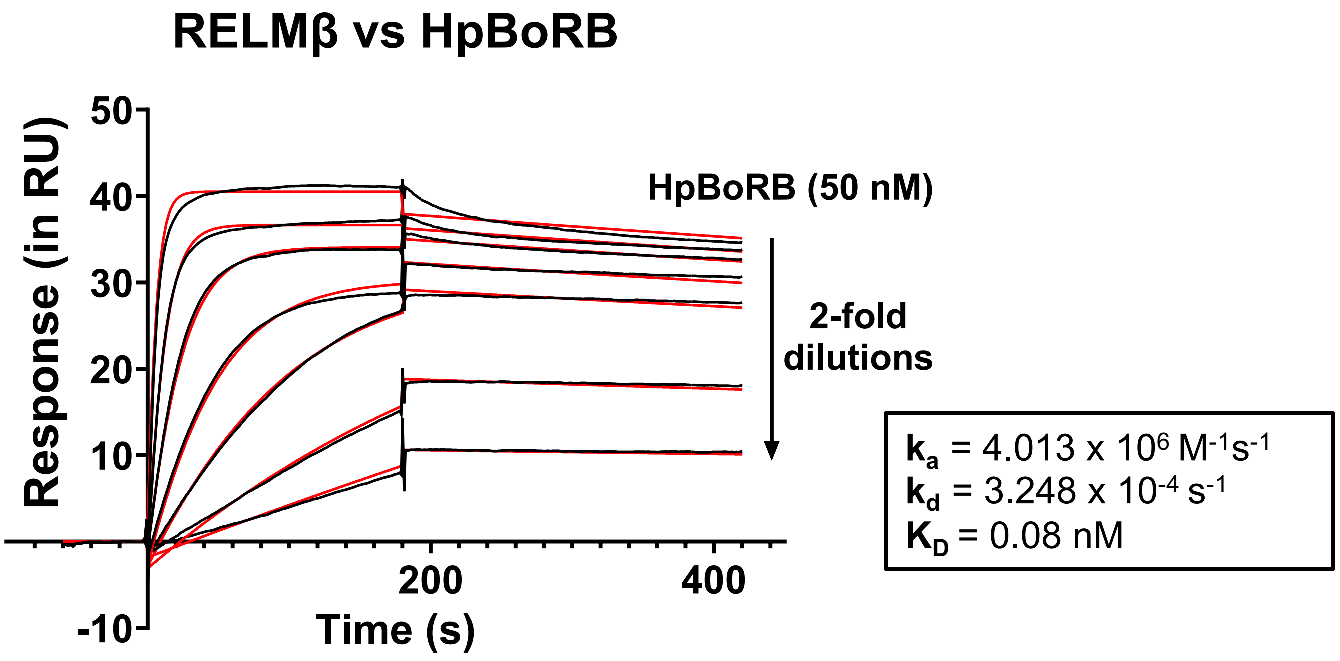


**Figure 6 - Surface plasmon resonance to determine HpBoRB - RELMβ interaction**
A surface plasmon resonance sensogram showing the binding of a concentration series (two-fold dilutions from 50 nM) of HpBoRB to immobilised murine RELMβ. Black lines show SPR data and red lines fit a 1:1 binding model. Data from a single experiment.

### Supplementary Figure 1

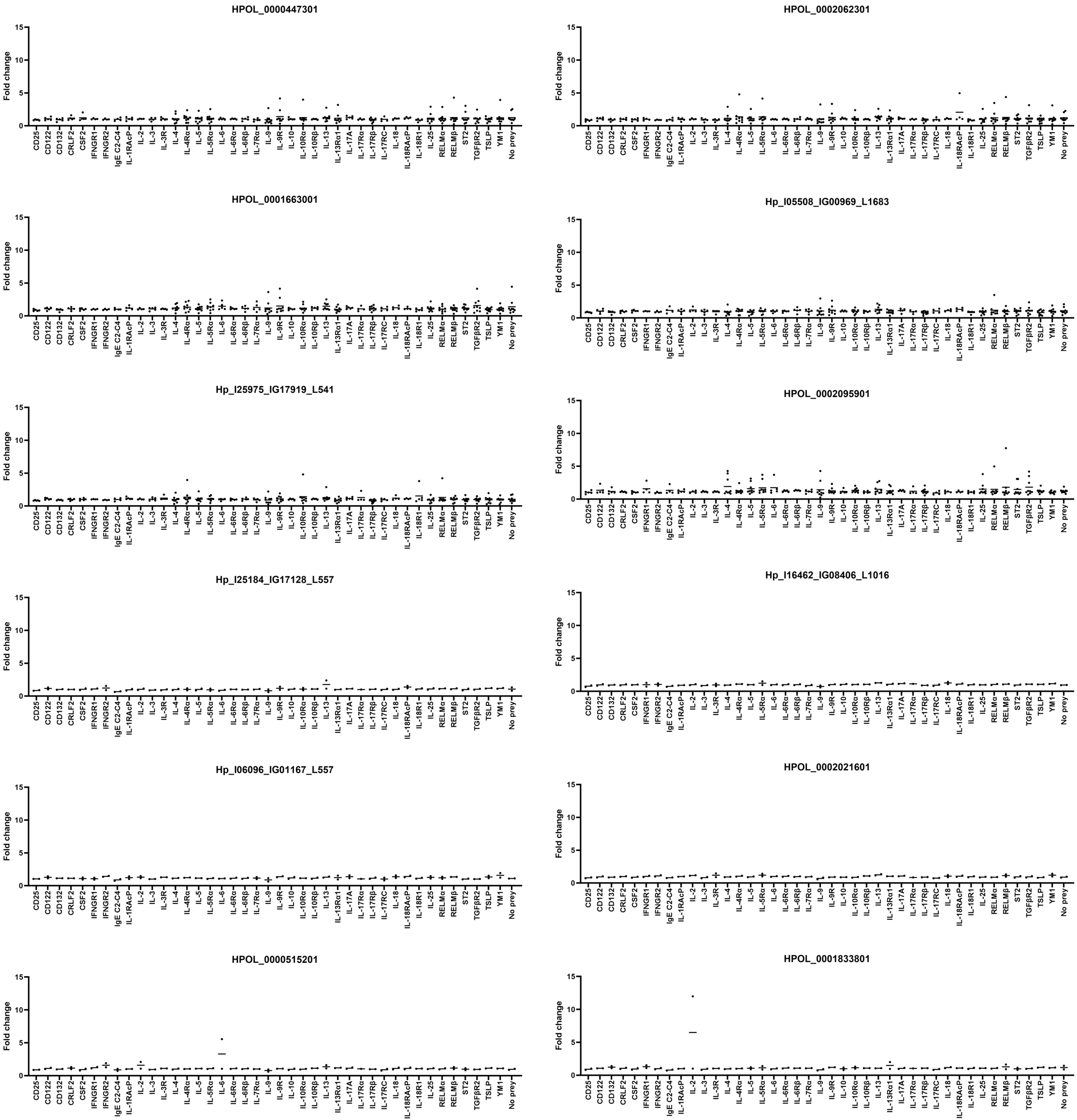

### Supplementary Figure 1

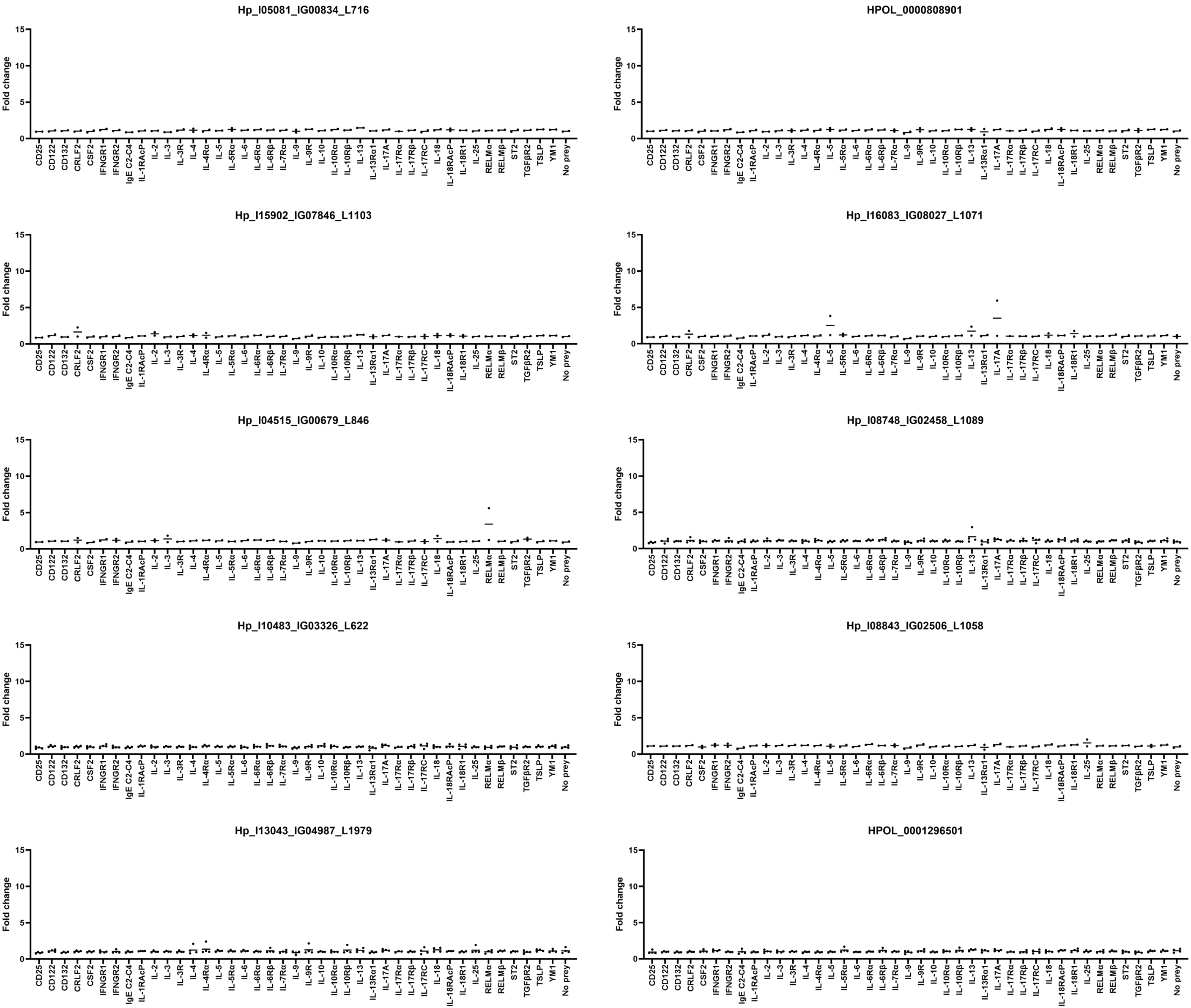


**Supplementary Figure 1 – Screening for protein-protein interactions**
All parasite baits shown in Figure 3 were screened against immune preys. Pooled data showing 2-10 replicates per condition, as indicated.

**A**

```
>HPOL_0001072601
ATGCTGCGAATCGTCACGTTCGCCATTCTAATCGTAACAGCAAAT
GCCAACTGTCCTGCCATCACACTTGATAAAGGCACAATAACGTAC
GACAAGGCTGCTGTCAATGGCCTGTATCCGGAAGGAACTTTCGCT
CATGGTCTGTGCCAGCAAGGCTTCAGCCTTTTCGGGCAATCGGGC
ACGAATTGCGGGAAAGACGGCAAATGGGATGGAGAACTTGGCAAA
TGCGATGTGAGCCCTCCGGTACAGGCATTCACCGAATGCTCGCCG
ACATATGTTCAAAACGGAGTGGTCACCTACGACCGAAACTCCGAT
CAGTTTTACAAACCAGAAGGAACGAAGGCAACCCTTACTTGCAAT
CAGGGATACAGACCAAGCGGCAACGCTACCGCTACGTGCGACAAA
GGAGAGTGGACTCCGGTTCTTGATCTTTGCGTTGTCTAA
```

**B**

MLRIVTFAILIVTANAN**C**PAITLDKGTITYDKAAVNGLYPEGTFA
HGL**C**QQGFSLFGQSGTN**C**GKDGK**W**DGELGK**C**DVSPPVQAFTE**C**SP
TYVQNGVVTYDRNSDQFYKPEGTKATLT**C**NQGYRPSGNATAT**C**DK
GE**W**TPVLDL**C**VV-

**C**

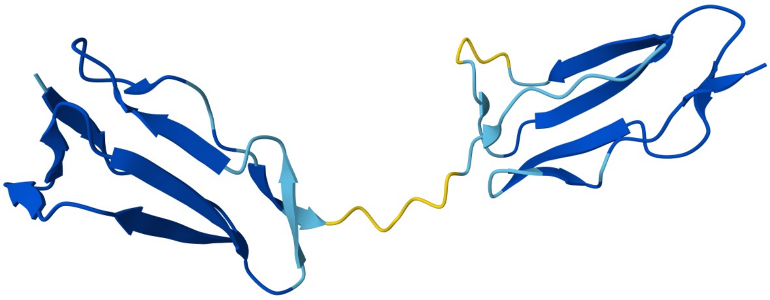

**Supplementary Figure 2 – Sequence and structure of HPOL_0001072601 (HpBoRB)**
DNA sequence of HPOL_0001072601 transcript shown in (A), with translated amino acid sequence in (B). In (B), predicted signal peptide highlighted in yellow, while 2 CCP domains are highlighted in grey, with characteristic Cys and Trp residues shown in bold. Alphafold3 model of HpBoRB sequence (minus signal peptide) shown in (C), with 2 CCP domains shown. Colour indicates Alphafold3 pIDDT confidence (yellow indicating low confidence, bleu indicating high confidence).

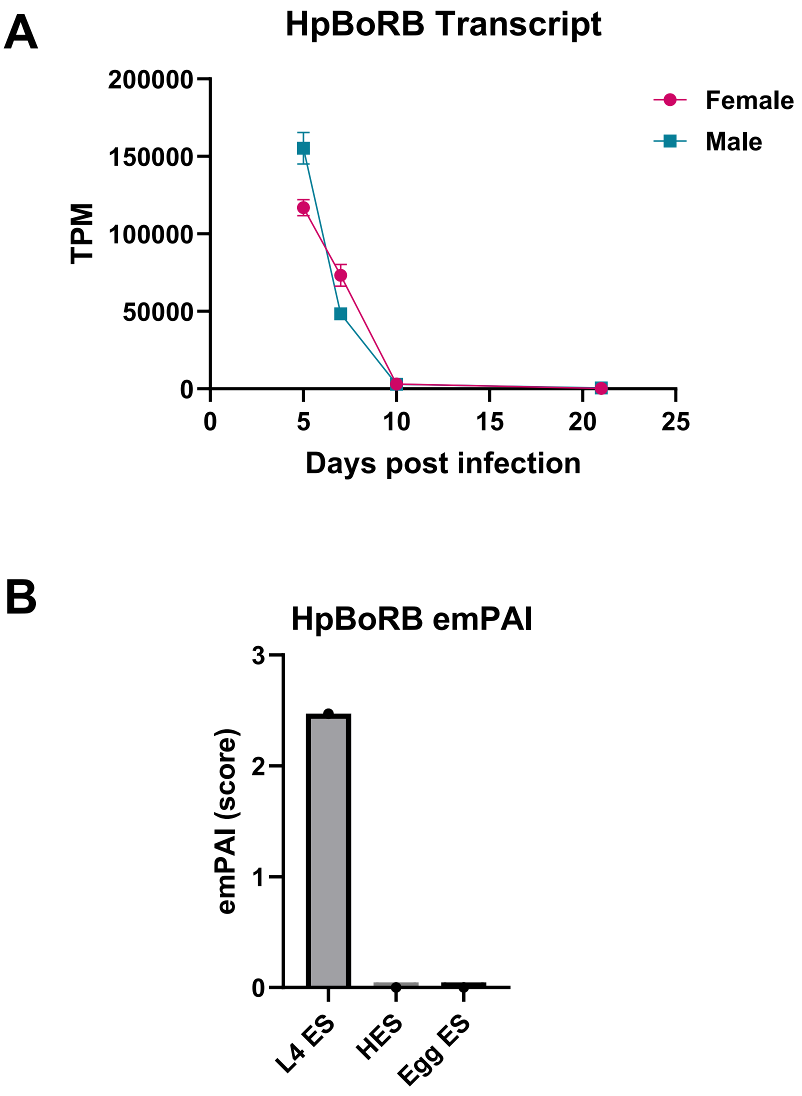


**Supplementary Figure 3 – Expression of HpBoRB during infection**
HpBoRB transcripts per million (TPM) in male and female *H. polygyrus bakeri* over a time course of infection was retrieved from Sequence Read Archive under accession number PRJNA750155 (A). HpBoRB protein expression in L4 ES, adult ES (HES) or egg ES was retrieved from Hewitson et al (B).
