## Supplementary table for "HpBoRB, a helminth-derived CCP domain protein which binds RELMβ"

**Supplementary Table 1: AVEXIS immune preys**

| **Target** | **Uniprot Accession** | **Region expressed** | **Notes** |
| --- | --- | --- | --- |
| CD25 | P01590 | 22-236 |  |
| CD122 | P16297 | 27-240 |  |
| CD132 | P34902 | 23-263 |  |
| CRLF2 | Q8CII9 | 20-232 |  |
| CSF2 | P26955 | 23-441 |  |
| IFNGR1 | P15261 | 26-254 | Variant:  rs13467528 Glu95Gly |
| IFNGR2 | Q63953 | 20-243 | Mutation:  Ser220Pro |
| IgE | P06336 | 91-421 | Heavy chain constant region |
| IL-1RAcP | Q61730 | 21-367 |  |
| IL-2 | P04351 | 21-169 |  |
| IL-3 | P01586 | 27-166 |  |
| IL-3R | P26952 | 17-331 | Variant:  rs52451947 Asp113Gly |
| IL-4 | P07750 | 21-140 |  |
| IL-4Rα | P16382 | 26-233 | Variant:  rs241116816 Cys59Arg  rs235962229 Met193Thr |
| IL-5 | P04401 | 21-133 |  |
| IL-5Rα | P21183 | 18-339 |  |
| IL-6 | P08505 | 25-211 |  |
| IL-6Rα | P22272 | 20-364 |  |
| IL-6Rβ | Q00560 | 23-617 |  |
| IL-7Rα | P16872 | 21-239 |  |
| IL-9 | P15247 | 19-144 |  |
| IL-9R | Q01114 | 38-270 | Insertion:  Glu192_Ala193insGln  Variant:  rs249160717Ile217Val |
| IL-10 | P18893 | 19-178 |  |
| IL-10Rα | Q61727 | 17-241 |  |
| IL-10Rβ | Q61190 | 20-220 |  |
| IL-13 | P20109 | 19-131 |  |
| IL-13Rα1 | O09030 | 26-340 |  |
| IL-17A | Q62386 | 26-158 |  |
| IL-17Rα | Q60943 | 32-322 |  |
| IL-17Rβ | Q9JIP3 | 18-286 |  |
| IL-17RC | Q8K4C2 | 22-464 |  |
| IL-18 | P70380 | 36-192 |  |
| IL-18RAcP | Q9Z2B1 | 20-356 |  |
| IL-18R1 | Q61098 | 20-326 |  |
| IL-25 | Q8VHH8 | 17-169 |  |
| RELMα | Q9EP95 | 24-111 |  |
| RELMβ | Q99P86 | 24-105 |  |
| ST2 | P14719 | 27-326 |  |
| TGFBR2-2 | Q62312-1 | 24-194 | Isoform RII-2 |
| TSLP | Q9JIE6 | 20-140 |  |
| YM1 | O35744 | 22-398 |  |
